# Patient-Derived Three-Dimensional Lung Tumor Models to Evaluate Response to Immunotherapy

**DOI:** 10.1101/2025.05.19.654890

**Authors:** Kayla F. Goliwas, Kenneth P. Hough, Sruti Sivan, Sierra L. Single, Sameer S. Deshmukh, Joel L. Berry, Maya Khalil, Benjamin Wei, Yanis Boumber, Mohammad Athar, Aakash Desai, Selvarangan Ponnazhagan, James M. Donahue, Jessy S. Deshane

## Abstract

Novel preclinical models that better mimic the *in vivo* tumor microenvironment are needed for advanced understanding of tumor biology and resistance/response to therapy. Herein, we report development of a novel *ex vivo* patient-derived three-dimensional lung tumor model (3D-LTM), that maintains features of human extracellular matrix, cell-cell interactions, and tissue architecture to evaluate a rapid response to immune checkpoint inhibitors (ICI). Within this model system, we recapitulated the heterogeneity of response to immunotherapy observed in non-small cell lung cancer (NSCLC) patients and defined signatures associated with response for predicting early response of ICI in patients. Spatial transcriptomics of the 3D-LTMs identified positive correlation of CD8^+^ T cell populations, CD4^+^ memory T cells, mast cells, NK cells, endothelial cells and non-classical monocytes with response status, whereas macrophages negatively correlated with response status. Pathway analysis of gene expression showed that chemokine signaling related pathways were activated in responder 3D-LTM tissues, whereas suppression of antigen presentation-related pathways and activation of T_reg_ differentiation-related pathways was associated with 3D-LTMs that were not considered responders. This model system has utility for rapid testing of novel immune directed therapy outcomes and for developing biomarkers of ICI response in NSCLC.

## INTRODUCTION

Lung cancer is the leading cause of cancer related death worldwide, with non-small cell lung cancer (NSCLC) being the most common subtype comprising 80-85% of lung cancer diagnoses (1). While therapeutic advances have been made, including novel molecular targeted therapies and immunotherapies, only a subset of patients receive long term benefit from these therapies and variable responses are common (2). This is in part due to the heterogeneity associated with the lung tumor microenvironment (TME) (3,4), which includes not only malignant tumor cells but also stromal cells, immune cells, extracellular matrix, and secreted factors including cytokines and chemokines. Both inter-and intra-patient variability in NSCLC TME leads to heterogeneity in therapeutic response (5). Yet, experimental models that represent this heterogeneity are lacking.

While model systems that accurately mimic patients’ tumors are crucial for translational cancer research, the highly utilized models do not recapitulate the true complexity of the lung TME. This includes tumor cell lines, which were originally derived from cancer patients and are the most commonly utilized *in vitro* model. The predictive value of cell culture models is limited, as cells are passaged in non-physiologic culture conditions that do not accurately mimic the tissue stiffness, oxygen or nutrient availability, or three-dimensional (3D) TME. Furthermore, propagation for multiple generations often modifies the genetic landscape and therapeutic responses observed (6–8). Xenografts tumors of these cell lines in immunodeficient mice are the most commonly used *in vivo* tumor models. While these models closely mimic the physiologic environment, they are still often not predictive of clinical response (9).

Patient-derived xenografts (PDX) have emerged as a useful alternative, as these models maintain tumor heterogeneity and preserve critical features of the original cancer for at least several passages (10–12). These models can replicate the complex cellular architecture of human tumors and can be valuable in drug development pipelines. However, PDX models of NSCLC are associated with limitations, including a 30-40% success rate, substantial time and resources to develop, and more importantly lack human immune system components as they are generally generated in NSG mice, limiting their utility in studying immunotherapy responses (12–16). The latest evolution in cancer modeling has been the development of patient-derived organoids (PDOs). These three-dimensional structures better recapitulate the histologic and genetic characteristics of patients’ tumors (17–19). Yet even these models face challenges; many organoid systems primarily represent tumor epithelium, while stromal and immune cells are excluded or gradually lost during *ex vivo* culture (20). Furthermore, generation of these models involve dissociation of the native TME as they require a high degree of tumor purity for success (20,21). The variable success rates in establishing these lung tumor models, and the potential overgrowth of normal airway cells present additional hurdles (22–24). Some promising PDO variations including the development of assembloid systems that incorporate multiple cell types and the use of microfluidic platforms to model cell-cell and cell-matrix interactions have been described (25,26), yet these systems still have their limitations, including lack of native human extracellular matrix (ECM) and tissue level dimensionality, and do not include all aspects of the human TME.

Thus, next-generation preclinical models that better integrate various components of the TME to more accurately mimic human tumors and patient responses are needed. Herein we describe an *ex vivo* 3D tissue culture platform that allows patient-derived NSCLC to be cultured via a perfusion bioreactor. This platform utilizes 5 mm tumor tissue cores, which maintain the native tissue architecture and tumor-stroma interactions over the culture period. Using this model, we have assessed response to commonly utilized immune checkpoint inhibitors (ICI) including a programmed death-ligand 1 (PD-L1) blocking antibody and a programmed death-1 (PD-1) blocking antibody to determine therapeutic response and changes in spatial immune cell landscape and transcriptomic signatures in response to ICIs. Using this platform, we demonstrate the heterogeneity of response to ICI observed in NSCLC patients and have delineated response groups and defined gene signatures associated with different response groups which could be useful for determining early response of ICI in patients.

## METHODS

### Clinical Sample Collection

De-identified, remnant surgical specimens were obtained from lobectomy and wedge resection surgeries performed on NSCLC patients at the University of Alabama at Birmingham Hospital. This study was approved by the University of Alabama at Birmingham Institutional Review Board (IRB-300003092, IRB-300003384, IRB-300008998) and conducted following approved guidelines and regulations Patient demographics are described in **Tables 1** and **2**. <u>Sex as a Biological Variable:</u> Tissue specimens from both males and female patients were used in this study and sex as a biological variable was considered.

### Sample Processing and *Ex Vivo* Perfusion Culture

Five mm diameter tissue cores were produced from remnant surgical tumor specimens using a tissue coring press (Alabama Research and Development, USA). As previously described (27), one tissue core was placed into the central chamber of each polydimethylsiloxane (PDMS, Krayden, USA) bioreactor within a liquid mixture of ECM components (90% collagen type 1 (Advanced Biomatrix, USA) + 10% growth factor reduced Matrigel (Corning, USA)) for structural support (28,29). The tissue/ECM volume was then perforated with five 400-micron Teflon coated stainless steel wires to generate through-channels for tissue perfusion. Following ECM polymerization, wires were removed, and the through-channels were filled with tissue culture media (1:1 mixture of X-Vivo15 and Bronchial Epithelial Growth media (Lonza, USA) with antibiotics (MP Biomedicals, USA)). Bioreactors were then connected to a perfusion system and tissue culture media was perfused through the tissue volume for 7 to 14 days (37°C, 5% CO_2_), with media changed every 3 days, as previously described for 3D lung tissue models (27). For therapeutic intervention experiments, immune checkpoint blocking antibody (10 μg/mL *InVivo*MAb anti-human PD-L1 (BioXCell, Catalog number: BE0285) or *InVivo*MAb anti-human PD-1 (BioXCell, catalog number: BE0188)) or matched IgG control antibody (BioXCell; *InVivo*MAb mouse IgG2b isotype control (catalog number: BE0086) or *InVivo*MAb mouse IgG1 isotype control (catalog number: BE0083), respectively) was introduced into perfusion culture starting on day 7 and treatment occurred every 3 days with media change. At the end of each experiment, the tissue specimen was removed from the bioreactor and split for (a) histologic processing and (b) collagenase B (Roche, Switzerland) digestion for flow cytometry analysis. Conditioned media (circulating tissue culture media) was collected with each media change for cytokine profiling and lactate dehydrogenase (LDH) measurement.

### Histologic Analysis

A portion of each starting tissue (day 0) and three-dimensional lung tumor model (3D-LTM) was fixed in 10% neutral buffered formalin for 24-48 hours prior to paraffin embedding. 5-micron histologic sections were subsequently generated from tissue blocks for hematoxylin and eosin (H&E) staining, Masson trichrome staining, immunostaining, and GeoMx Digital Spatial Profiling (DSP). <u>Cell Density Analysis</u>: Sections were H&E stained, and photomicrographs were acquired to evaluate tissue architecture and cell density (number of cells per cross-sectional area). <u>Immunohistochemistry (IHC):</u> Following antigen retrieval (BioGenex Citra Plus, pH 6 (catalog number: HK086-5K)), IHC was performed to detect cleaved caspase 3 expression (1:100, clone D3E9, Cell Signaling) using the Leica Novolink Polymer Detection System (catalog number: RE7290-CE).

### Matrix Proteomics

For matrix protein enrichment and extraction, tissue samples were prepared as described previously (30,31). Briefly, tissues were processed using the Millipore Compartment Protein Extraction Kit with some modifications of the described methodology(30) and all fractions were stored at −80° overnight. The ECM fraction was then reconstituted in 8M urea and deglycosylated. The urea-insoluble fraction was collected by centrifugation, reconstituted in 1x LDS sample buffer, and sonicated for 20 minutes in an ultrasonic water bath. Both urea-soluble and insoluble fractions were quantified via EZQ protein assay, and an equal amount per sample was loaded onto 10% Bis-tris gels and gels were stained overnight with Colloidal Coomassie. Each sample was then digested in 3 fractions with trypsin overnight and high-resolution LC-ESI-MS/MS analysis was completed. Data was searched against the human subset of the Uniref100 database with Carbamidomethylation, Oxidation, and Hydroxyproline.

### Cell Viability Analysis

<u>Flow Cytometry Analysis:</u> Following tissue digestion, single cell suspensions were stained with the BD Pharmingen PE Annexin V Apoptosis Detection Kit I (catalog number: 559763) following manufacturer’s instructions. Analyses were performed on FACSymphony A3 Cell Analyzer with FACSDiva software version 8.0.1 (BD Biosciences, Germany). Data were analyzed with FlowJo 10.7.1 (Treestar, USA). <u>TUNEL Staining:</u> TUNEL staining was completed using the TumorTACS In Situ Apoptosis Detection Kit (R&D Systems; catalog number: 4815-30-K) as per manufacturer instructions. <u>Measuring LDH</u>: LDH was measured in conditioned media using the Invitrogen CyQUANT LDH Cytotoxicity Assay (Thermo Fisher, Germany) following manufacturer’s instruction.

### Multiparameter Flow Cytometry

The following antibodies were used for multiparameter flow cytometry: Anti-CD3-alexafluor 700 (Clone: UCHT1); anti-CD11b-APC-Cy7 (Clone: ICRF44); anti-CD90-BV421 (Clone: 5E10) from BD Biosciences (Germany). Anti-HLA-DR-APC (Clone: LN3); anti-CD3-PE-Cy7 (Clone: UCHT1); anti-CD8-APC (Clone: OKT8); anti-CD64-PerCp-eFluor710 (Clone: 10.1); anti-CD14-PE/Cyanine 7 (Clone: 61D3); anti-CD163-alexafluor 700 (Clone: GHI/61) from eBioscience (Thermo Fisher, Germany). Anti-Ki-67-Dylight350 (Clone: 1297A); anti-Carbonic Anhydrase IX-PE-Cy7 (Clone: 2D3) from Novus (USA). Anti-CD45-APC-Cy7 (Clone: 2D1); anti-PD-1-BV605 (Clone: NAT105); anti-CD16-PE (Clone: 3G8); anti-CD45-Pacific Blue (Clone: HI30); anti-CD66b-PerCP-Cy5.5 (Clone: G10F5); anti-EpCAM(CD326)- Alexafluor 594 (Clone: 9C4); anti-CD31-Alexafluor 700 (Clone: WM59); anti-PD-L1(CD274)- PerCP-Cy5.5 (Clone: 29E.2A3); anti-CD15-Alexafluor 700 (Clone: W6D3) from Biolegend (USA). The Foxp3 / Transcription Factor Staining Buffer Set (Thermo Fisher, Germany) and the Cytofix/Cytoperm Fixation/Permeablization kit (BD, Germany) were used according to the manufacturer’s protocol to stain for intracellular molecules (intranuclear and cytoplasmic molecules, respectively). Analyses were performed on FACSymphony A3 Cell Analyzer with FACSDiva software version 8.0.1 (BD Biosciences, Germany). Data were analyzed with FlowJo (Treestar, USA).

### Multiplex Cytokine Analysis

Soluble cytokines were evaluated in conditioned media from 3D-LTMs using a Bio-Plex200 and the Bio-Plex Pro Human Cytokine 17-plex Assay (catalog number: M5000031YV) following manufacturer’s protocol.

### GeoMx Digital Spatial Profiling

Spatial transcriptomic profiling was completed on histologic sections of αPD-1, αPD-L1, and IgG treated 3D-LTMs using the GeoMx Digital Spatial Profiler (DSP) with the GeoMx Human Cancer Transcriptome Atlas (Nanostring Technologies). Briefly, 5 μm FFPE sections were stained with morphology markers by first incubating slides overnight at 38° C followed by an additional 2 hours at 60° C. Then sections were manually stained with the Human Cancer Transcriptome Atlas UV-cleavable barcoded RNA probes along with antibodies against human Pan-cytokeratin, CD45, and CD8 as well as SYTO 13 for geometric region of interest (ROI) selection. Using the GeoMx DSP, ROIs were selected, and barcoded RNA probes were cleaved and collected from each ROI. Library Prep with Seq-Code primers was performed and libraries were sequenced on an Illumina sequencing instrument. FASTQ files were then converted into digital count conversion (DCC) files using GeoMx NGS Pipeline and uploaded onto the GeoMx Analysis Platform. Data underwent quality control and Q3 normalization prior to analysis.

#### Bioinformatics and Statistical Analysis

Spatial transcriptomics data were obtained from NanoString GeoMx DSP outputs, including raw expression counts, segment metadata, and probe annotations. Gene identifiers and segment display names were standardized using make.names() to ensure consistency across datasets. Negative control probes were identified based on the presence of “Neg” in the target name, and background signal was estimated using the derive_GeoMx_background() function from the SpatialDecon package (v1.4.0). Signal-to-noise ratios (SNR) were calculated by dividing raw counts by background estimates. A background-subtracted SNR matrix was further processed using quantile normalization, scaling each segment by the 85th percentile of its expression distribution. Region of interest (ROI) filtering was applied by computing the 85th percentile signal across ROIs; segments with signal values below 1.0 were excluded from downstream analysis. Normalization strategies tested included background subtraction and scaling. Clustered heatmaps with dendrograms were generated and unpaired Student’s t-tests with Benjamini-Hochberg correction were performed to compare ICI treatment to matched IgG controls. Ultimately, raw SNR values (background-normalized) were selected for deconvolution. ROI-level data were further aggregated into ROI-level matrices stratified by ROI type (e.g., tumor, stroma). For each ROI type, matrices were constructed by matching ROI identifiers with corresponding segment names. Genes were considered suitable for analysis if they exhibited sufficient variation across samples (standard deviation > 0.2 and maximum expression > 2.5). These thresholds were applied both globally and per ROI type to ensure robustness of downstream modeling. Cell type proportions were estimated using the SpatialDecon package, utilizing the spatialdecon() function. A reference profile matrix (safeTME and lung+neutrophil) was used as the basis for cell-type-specific expression patterns (32). Deconvolution was performed with 1000 iterations, incorporating cell count estimates from each ROI (segment-level nuclei counts) as weights. Background-corrected abundance scores (beta) estimated proportions of non-tumor content, and inferred cell counts were retained for further analysis and export. Floret plots were generated to visualize cell-type compositions within each ROI. Each plot depicted a polygonal boundary inferred from ROI coordinates and a radial “floret” indicating relative cell-type contributions, scaled by square root transformation for visual clarity. Data processing and statistical analysis were conducted using custom R scripts (version 4.3.2 ("Eye Holes") on a 64-bit Linux platform (x86_64-pc-linux-gnu)) with several packages from the Bioconductor and CRAN ecosystems. <u>Differential Gene Expression</u>: Raw gene expression counts were imported from a tab-delimited file containing count data. Non-integer values were rounded to the nearest integer to ensure compatibility with the DESeq2 framework. Sample metadata was manually curated into a design matrix specifying treatment conditions, including anti-PD-1, anti-PD-L1, and their respective isotype controls (IgG1 and IgG2B), and treatment response types. Differential gene expression analysis was performed using the DESeq2 R package (v1.40.2). A DESeq2 dataset was constructed using DESeqDataSetFromMatrix() with ∼ condition as the design formula. The DESeq2 pipeline was executed via the DESeq() function, and contrasts between treatment conditions were extracted using the results() function. Volcano Plots: Volcano plots were generated using ggplot2 (v3.4.4) with custom annotations highlighting the top 10 most significant genes based on adjusted p-value (FDR). Genes were categorized as significant if they met both criteria: |log₂ fold change| ≥ 1 and adjusted p-value (padj) < 0.05. Non-significant genes and those with missing p-values were appropriately adjusted or excluded from the visualizations. <u>Heatmaps</u>: Curated gene sets representing biologically relevant pathways, such as Tumor Inflammation Signature, T Cell Checkpoints, Chemokine Signaling, MHC Class II Antigen Presentation, Dendritic Cell Activation, and Mast Cells & IL-3 Signaling, were defined manually based on known functional roles defined by NanoString. For pathway-level summaries, one-way ANOVA was performed per gene across treatment groups to identify the top 20 differentially expressed genes within each pathway. For each treatment group, two sample *t*-tests were used to compare treated and control conditions, and gene-level summary statistics, including means, log2 fold-change, and adjusted *p*-values, were compiled. Results were visualized as log2 fold-change heatmaps using the ComplexHeatmap package, clustered by gene where appropriate. <u>Pathway Analyses</u>: To identify enriched biological processes, two complementary gene ontology (GO) enrichment approaches were performed using the clusterProfiler (v4.10.0), org.Hs.eg.db (v3.18.0), and DOSE packages. Overrepresentation analysis (ORA) was conducted using enrichGO() for genes that were significantly differentially expressed (|log₂ fold change| ≥ 1, padj < 0.05). Analyses were performed across three GO domains: Biological Process (BP), Molecular Function (MF), and Cellular Component (CC), with the option to also aggregate all terms (ALL). Gene set enrichment analysis (GSEA) was performed using gseGO() with log₂ fold change values ranked in descending order. GSEA parameters included a minimum gene set size of 3, maximum of 500, and 10,000 permutations (nPermSimple = 10000). Pathway visualizations were generated using dotplot() and gseaplot2() from enrichplot (v1.20.0), and top pathways were reported. Statistical analyses were performed using GraphPad Prism software (La Jolla, CA, USA), the GeoMx Analysis Platform, or R. Data are presented as mean ± standard deviation unless indicated otherwise in the figure legend. A p-value less than 0.05 was considered statistically significant. A two-tailed unpaired Student’s t-test was performed to compare the two groups. One-way ANOVA with Tukey’s multiple comparisons test or Sidak’s multiple comparison test was performed to compare data with more than two groups to determine statistical significance.

## RESULTS

### Perfused Explant 3D Lung Tumor Models Retain Tissue Architecture, Extracellular Matrix and Cell Viability

Explant 3D lung tumor models (3D-LTMs) from patient lung tumors were generated using a perfusion bioreactor system previously utilized for lung tissue exposure and 3D cell culture studies (27–29,31,33). As shown in **Fig. 1A**, remnant NSCLC tissues from treatment naïve patients undergoing tumor resection (patient demographics described in **Supplemental Table 1**) were obtained through IRB approved protocols and 5 mm tissue cores were generated using a tissue coring press. One tissue core was placed within a liquid ECM mixture in our fabricated silicon bioreactor and 5 teflon-coated wires penetrated the tissue/ECM volume to generate through-channels for media perfusion. Following ECM polymerization, wires were removed, tissue culture media was added to perfusion channels, and the bioreactor was connected to a perfusion loop. Following 7- or 14-day *ex vivo* culture, tissue was removed and split for histologic analysis, cell phenotyping via flow cytometry, and matrix proteomics. When histologic cross-sections were assessed, cell density (average number of cells per area) was maintained in 3D-LTMs cultured for 7 and 14 days when compared to starting tissue (D0, **Fig. 1B**). To ensure that the human ECM was being maintained in 3D-LTMs, proteomics analysis was completed. This analysis showed the relative proportion of most ECM protein components was maintained in 3D-LTMs cultured for 14 days when compared to starting tissue, with some loss in proteoglycans (**Fig. 1C**). Histologic architecture was monitored in H&E and Masson’s Trichrome stained cross-sections of starting tissue (left) and 3D-LTM tissue following 7 (middle) and 14 days perfusion (right, **Fig. 1D**) and found to be similar across tissues. Cell viability assessed by Annexin V/7AAD, **Fig. 1E-F**), cleaved caspase 3, (**Fig. 1G**), or circulating lactate dehydrogenase levels (LDH, **Fig. 1H**) did not differ between starting tissue (D0) and 3D-LTMs cultured for 14 days (D14).

**Figure 1:**
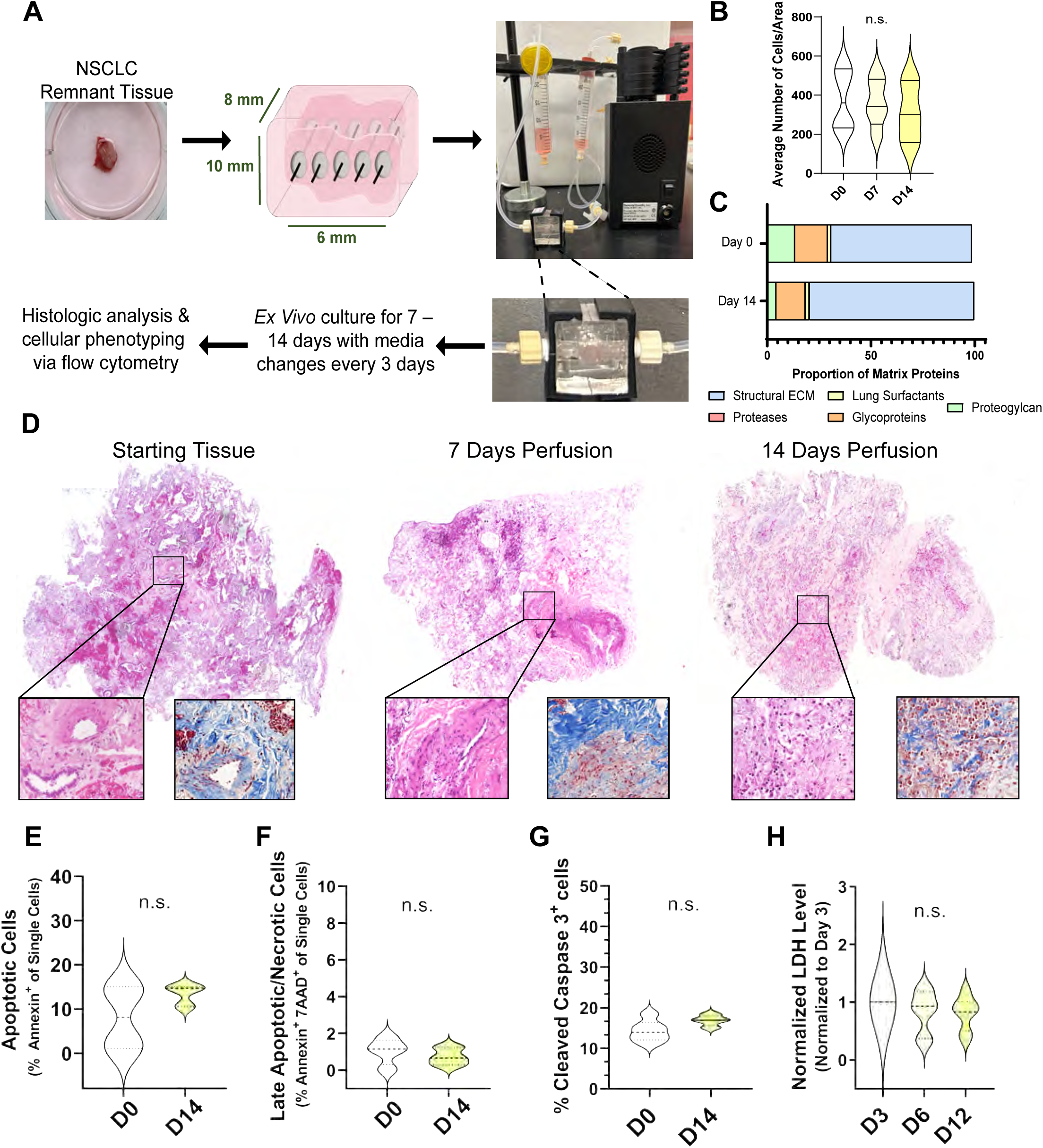
3D Lung Tumor Models Maintain Tissue Architecture, Extracellular Matrix and Cell Viability with Ex Vivo Perfusion Culture. **A.** Schematic of set up procedure for a 3D lung tumor model (3D-LTM) generated from patient lung tumors. **B.** Cell density (average number of cells per area) within histologic sections is maintained in 3D-LTM cultured for 7 and 14 days when compared to starting tissue (D0). **C.** Proteomics analysis shows the proportion of extracellular matrix protein classes is maintained in 3D-LTM cultured for 14 days when compared to starting tissue. **D.** Histologic cross sections of starting tissue (left) and 3D-LTM tissue following 7 (middle) and 14 days perfusion (right). **E-H.** Cell viability does not differ between starting tissue (D0) and 3D-LTM cultured for 14 days (D14).

We assessed the perfusion method to ensure appropriate maintenance of cell populations with *ex vivo* culture. Two perfusion methods were evaluated to assess optimal nutrient delivery; single pass media perfusion in which tissue culture media passed through the bioreactor once and was collected in a downstream collection reservoir, and recirculating perfusion in which tissue culture media circulated through the bioreactor and perfusion loop for 3 days before replacement. No change in the proportion of tumor cells, PD-L1^+^ tumor cells, or immune cells was observed in response to perfusion method (**Supplemental Figs. 1 & 2**). However, the proportion of endothelial cells significantly increased in 3D-LTMs cultured for 14 days using single pass media perfusion and fibroblast outgrowth was observed in 3D-LTMs cultured for 14 days using recirculating perfusion (**Supplemental Figs. 1 & 2**, * p<0.05). Therefore, single pass media perfusion was chosen for all additional 3D-LTM culture.

**Figure 2:**
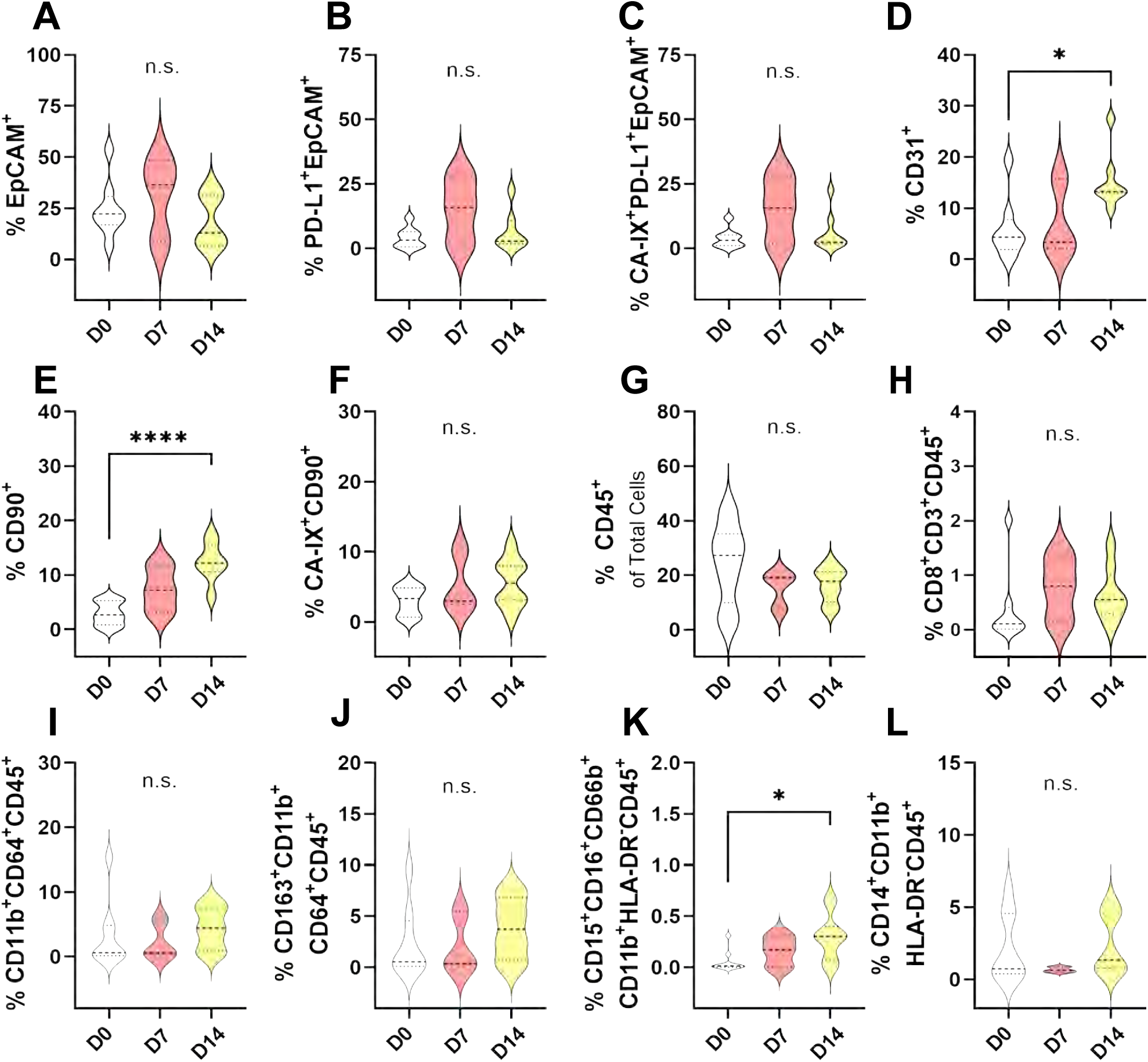
Immune and Non-Immune Cells within the Tumor Microenvironment are Preserved in 3D-LTM with Perfusion Culture. **A-C**. Proportion of tumor cells (**A**), PD-L1+ tumor cells (**B**), and hypoxic PD-L1+ tumor cells (**C**) in 3D-LTM following culture for 7 or 14 days when compared to starting tissue (D0). **D-F**. Shifts in the proportion of endothelial cells (**D**), fibroblasts (**E**), and hypoxic fibroblasts (**F**) in 3D-LTM following 14 days culture when compared to starting tissue (D0). **G-L**. Comparison of the proportion of immune cell populations in 3D-LTM following culture, including total immune cells (**G**), CD8+ T cells (**H**), macrophages (**I**) M2-like macrophages (**J**), granulocytic myeloid-derived suppressor cells (MDSC, **K**) and monocytic MDSC (**L**). * p<0.05; **** p<0.001.

### 3D Lung Tumor Models Maintain Tumor, Immune, and Stromal Cells with Perfusion Culture

Once single pass media perfusion was found to be the most effective perfusion method, cell phenotyping within 3D-LTMs was completed to ensure that cellular components remained similar to starting tissue (patient demographics found in **Table 1**). No change in tumor cells, PD-L1^+^ tumor cells, or hypoxic PD-L1^+^ tumor cells was observed in 3D-LTMs with culture. (**Fig. 2A-C**). While a significant increase in endothelial cells and fibroblasts was observed in 3D-LTMs following 14 days culture (**Fig. 2D-E**), no change in hypoxic fibroblasts or immune cell subsets, including CD8^+^ T cells, macrophages, M2-like macrophages, or monocytic myeloid-derived suppressor cells (MDSC), was noted in 3D-LTMs with culture (**Fig. 2F-J, 2L,** and **Supplemental Fig. 3**). The only immune cell population that differed in 3D-LTMs following culture was polymorphonuclear (PMN)-MDSC (**Fig. 2K**), however this cell type comprised a small proportion of the total cells within the tissue.

**Table 1:** Patient Demographics for Optimization of 3D-LTM.

| Case | Age | Sex | Race | Smoking Status | Histologic Subtype | Clinical Stage | TNM Stage | PD-L1 (Clinical IHC) | NGS Results/Mutation Status |
| --- | --- | --- | --- | --- | --- | --- | --- | --- | --- |
| 1 | 86 | M | White | Never | Adenocarcinoma | IIA | pT2bN0 | 0% | Kras G12C |
| 2 | 70 | M | White | Current | SCC | IB | pT2aN0 | Not Assessed | Not Assessed |
| 3 | 64 | M | White | Current | SCC | IA | pT1cN0 | 65% | Not Assessed |
| 4 | 68 | M | White | Current | Adenocarcinoma | IB | pT1bN0 | Not Assessed | Not Assessed |
| 5 | 69 | F | White | Current | Adenocarcinoma | IA3 | pT1cN0 | 5% | KRAS p.G12A |
| 6 | 74 | F | Black | Never | Adenocarcinoma | IIA | pT1cN1 | 20% | EGFR ex19 del |
| 7 | 56 | F | White | Never | Adenocarcinoma | IA3 | pT1cN0 | 0% | EGFR missense mutation p.L858R in exon 21 |
| 8 | 75 | F | White | Former | Adenocarcinoma | IA2 | pT1bN0 | <1% | Not Assessed |
| 9 | 61 | F | Asian | Current | Adenocarcinoma | IIB | pT3N0 | 0% | KRAS missense pG12C |
| 10 | 74 | F | White | Never | Adenocarcinoma | IIB | pT3N0 | 0% | Not Assessed |
| 11 | 60 | F | White | Former | Adenocarcinoma | IIB | pT4N0 | 0% | Not Assessed |
| 12 | 62 | F | White | Current | Adenocarcinoma | IA2 | pT1bN0 | 0% | Not Assessed |

### Patient-Derived 3D Lung Tumor Models Simulate *In Vivo* Response to αPD-L1

As 3D-LTMs were found to maintain the native tissue cell composition, ECM, and tissue architecture, we next determined whether these models could be utilized for therapeutic testing, focusing on immune directed intervention. As shown in **Figure 3A**, 3D-LTMs from 10 patient tumors (patient demographics found in **Table 2**) were generated and maintained with perfusion culture. Starting on day-7 of culture, 10 μg/mL anti-human PD-L1 blocking antibody or matched isotype control antibody (mouse IgG2b isotype control) was introduced into perfusion. Intervention subsequently occurred every 3 days with media change until the *ex vivo* culture was stopped on day-14. 3D-LTMs were categorized as responders, partial responders, or non-responders based on changes in the ratio of PD-L1^+^ tumor cells to total tumors cells and the ratio of PD-1^+^ CD8^+^ T cells to total CD8^+^ T cells. 3D-LTMs were considered responders if the difference in the ratio of PD-L1^+^ tumor cells to total tumors cells between patient-matched IgG and αPD-L1 treated 3D-LTM was below −0.1 and the difference in the ratio of PD-1^+^ CD8^+^ T cells to total CD8^+^ T cells was negative. 3D-LTMs were partial responders if the difference in the ratio of PD-L1^+^ tumor cells to total tumors cells was negative but not below the threshold of −0.1, and non-responders if the difference in the ratio of PD-L1^+^ tumor cells to total tumors cells was positive. Changes in total cell viability were observed in responder (green), partial responder (purple) and non-responder (red) 3D-LTMs (**Fig. 3B-D**). The proportion of total tumor cells within 3D-LTMs did not significantly change with αPD-L1 antibody treatment when all samples were combined (**Supplemental Fig. 4A**, left). However, when samples were split by responder status, a reduction in this cell population was observed in responder 3D-LTMs, as well as some partial and non-responder 3D-LTMs treated with αPD-L1 (**Supplemental Fig. 4A**, right). The overall % PD-L1^+^ tumor cells showed a decreasing trend in αPD-L1 antibody treated 3D-LTMs when compared to IgG controls (**Fig. 3E**, left), with a substantial reduction in % PD-L1^+^ tumor cells noted in responders (2 of 2 samples, 0.36 fold when compared to matched IgG, not significant due to limitation in number of samples) and partial responder 3D-LTMs (3 of 5 samples, 0.83 fold when compared to matched IgG; IgG vs. αPD-L1, t-test: p=0.12), but not the non-responder 3D-LTMs (**Fig. 3E**, right). However, when αPD-L1 treated partial responder 3D-LTM were compared to αPD-L1 treated non responders 3D-LTM a significant increase in the proportion of PD-L1^+^ tumor cells was observed in non-responder samples (p<0.0001).

**Figure 3:**
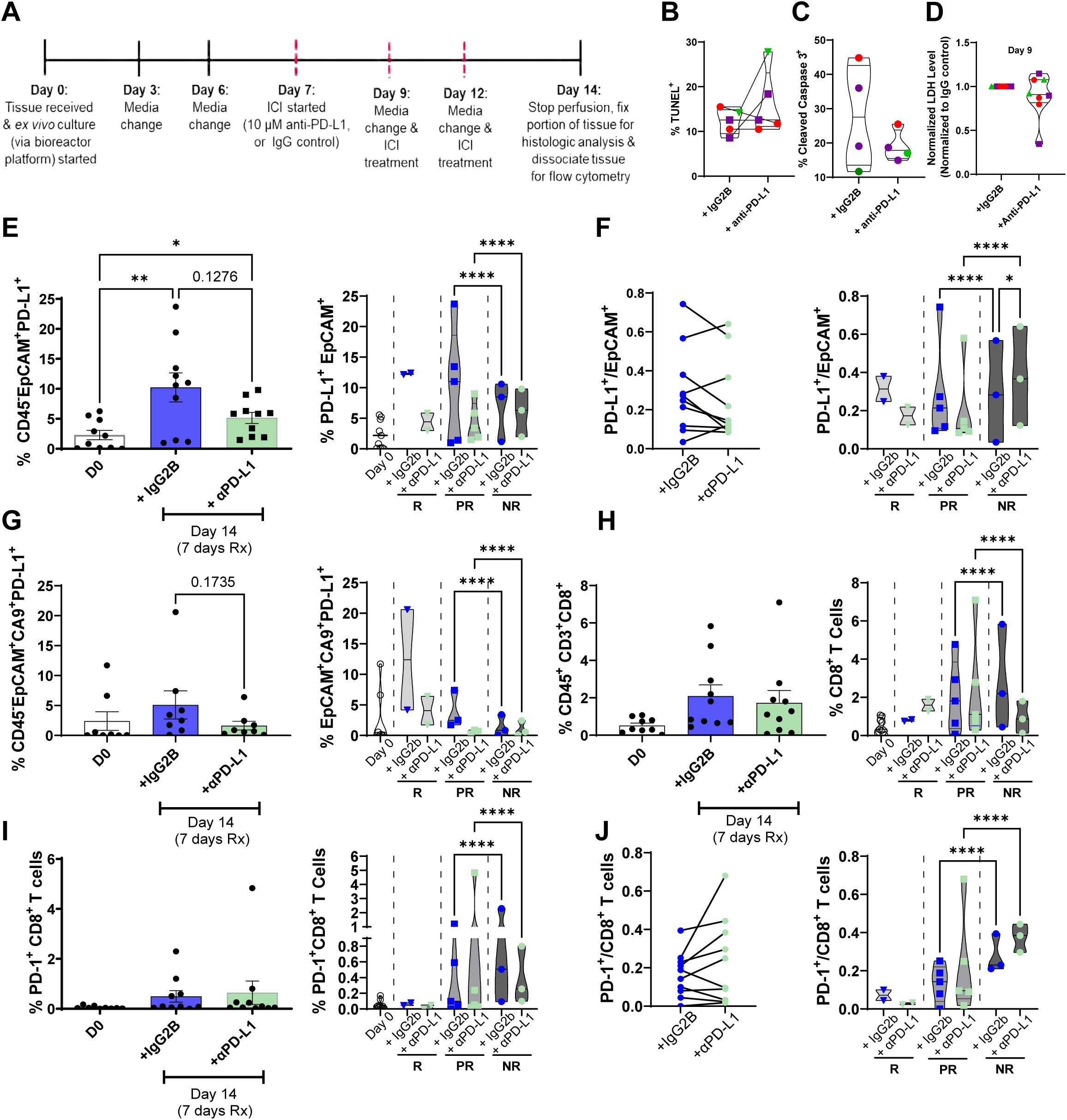
Patient-Derived 3D-LTM Model Response to αPD-L1. **A**. 3D-LTM αPD-L1 treatment schematic. **B-D**. Changes in cell viability are observed in responder (green), partial responder (purple) and non-responder (red) 3D-LTM. **E**. Shifts in the proportion of PD-L1 ^+^ tumor cells in 3D-LTM following αPD-L1 or IgG treatment (left), with data split by response status (right). **F**. The ratio of PD-L1 ^+^ tumor cells to total tumor used to determine response status (left), with samples are split by response status (right). **G-I**. Proportion of hypoxic PD-L1 ^+^ tumor cells (**G**), CD8^+^ T cells (**H**) and PD-1^+^ CD8^+^ T cells (**I**) in 3D-LTM following αPD-L1 or IgG treatment. **J**. The ratio of PD-1^+^ CD8^+^ T cells to total CD8^+^ T cells used to determine response status (left), with samples are split by response status (right). * p<0.05; **** p<0.001.

**Table 2:** Patient Demographics for 3D-LTM ICI Treatment.

| Case | Age | Sex | Race | Smoking Status | Histologic Subtype | Clinical Stage | TNM Stage | PD-L1 (Clinical IHC) | NGS Results/Mutation Status | 3D-LTM Blocking Antibody |
| --- | --- | --- | --- | --- | --- | --- | --- | --- | --- | --- |
| 1 | 78 | M | White | Current | Adenocarcinoma | IA2 | pT1bN0 | 0% | EGFR ex19 deletion | αPD-L1 |
| 2 | 74 | M | White | Former | Adenocarcinoma | IA2 | pT1bN0 | 0% | EGFR p.L858R (exon 21) and p.S768I (exon 20) | αPD-L1 |
| 3 | 75 | F | White | Former | Adenocarcinoma | IA3 | pT1cN0 | 1% | Not Assessed | αPD-L1 |
| 4 | 72 | F | White | Never | Adenocarcinoma | IA3 | pT1cN0 | 1% | EGFR and TP53 | αPD-L1 |
| 5 | 75 | M | White | Current | Adenocarcinoma | IA3 | pT1cN0 | 0% | KRAS missense p.G12C | αPD-L1 |
| 6 | 74 | M | White | Never | SCC | IIIA | pT4N1 | 80% | Not Assessed | αPD-1 |
| 7 | 67 | F | Asian | Former | SCC | IB | pT2aN0 | 1% | Not Assessed | αPD-1 |
| 8 | 73 | F | Black | Never | Adenocarcinoma | IA3 | pT1cNo | 0% | Tissue not sufficient | αPD-1 |
| 9 | 72 | F | White | Never | Adenocarcinoma | Ib | pT2aN0 | 0% | Not Assessed | αPD-L1/<br>αPD-1 |
| 10 | 67 | F | Black | Former | Adenocarcinoma | IIIA | pT2N2 | 30% | Tissue not sufficient | αPD-1 |
| 11 | 51 | M | White | Current | SCC | IB | pT3N0 | 0% | Not Assessed | αPD-L1 |
| 12 | 60 | M | White | Former | Adenocarcinoma | IIB | pT3N0 | 0% | Not Assessed | αPD-L1/<br>αPD-1 |
| 13 | 62 | M | White | Former | Adenocarcinoma | IIB | pT2aN1 | 10% | No significant mutation found | αPD-L1 |
| 14 | 62 | M | White | Former | SCC | IIIA | pT3N1 | 60% | PIK3CA p.E545Q c.1633G>C | αPD-L1/<br>αPD-1 |

70% of 3D-LTMs had a decreased ratio of PD-L1^+^ tumor cells to total tumor cells indicating response or partial response to αPD-L1 treatment (**Fig. 3F**, left); a reduction in this ratio was observed in all responder and partial responder 3D-LTMs, but a significant increase was observed in non-responder 3D-LTMs (p=0.02 (3 of 3 samples); **Fig. 3F**, right). Hypoxic PD-L1 ^+^ tumor cells also tended to decrease in αPD-L1 treated 3D-LTMs when compared to IgG control (**Fig. 3G**, left), with a reduction in hypoxic PD-L1 ^+^ tumor cells observed in responder and partial responder (IgG vs. αPD-L1, t-test: p=0.19) 3D-LTMs but not non-responder 3D-LTMs. Additionally, when αPD-L1 treated partial responder 3D-LTMs were compared to αPD-L1 treated non responder 3D-LTMs, a significant increase in the proportion of hypoxic PD-L1^+^ tumor cells was observed in non-responder samples (p<0.0001; **Fig. 3G**, right).

The overall proportions of total immune cells, CD8^+^ T cells, and PD-1^+^ CD8^+^ T cells were not altered with treatment (**Supplemental Fig. 4B** and **Fig. 3H-I**, left). However, total immune cells increased in responder 3D-LTMs and CD8^+^ T cells increased in responder (2 of 2 samples) and some partial responder (2 of 5 samples) 3D-LTMs, with this population of cells decreasing in non-responder 3D-LTMs (3 of 3 samples; **Supplemental Fig. 4B** and **Fig. 3H**, right). The proportion of PD-1^+^ CD8^+^ T cells was lower in responder 3D-LTMs compared to the other groups (**Fig. 3I**, left). Furthermore 30% of treated 3D-LTMs had a decreased ratio of PD-1^+^ CD8^+^ T cells to total CD8^+^ T cells in response to treatment (**Fig. 3J**, left), including both responder samples and 20% of the partial responder 3D-LTMs, whereas an increase in this ratio was observed in all non-responder 3D-LTMs (IgG vs. αPD-L1, t-test: p=0.13; **Fig. 3J**, right). Additionally, significant changes in the proportion of these cell populations were observed when IgG treated partial responder 3D-LTM were compared to IgG treated non responder 3D-LTM, suggesting baseline differences in these tissues.

Other cells within the TME were also evaluated following intervention. The overall proportions of G-MDSC, M-MDSC, fibroblasts, PD-L1^+^ fibroblasts, and endothelial cells (**Supplemental Fig. 4C-G**) did not change with αPD-L1 intervention (left). The proportions of both G-MDSC and M-MDSC, as well as endothelial cells were lower in responder 3D-LTMs (**Supplemental Fig. 4C-D** and **2G**, right). Additionally, PD-L1^+^ fibroblasts were reduced in responder and partial responder 3D-LTMs (**Supplemental Fig. 4F**, right). Together these data show a shift in the TME associated with response to treatment, with a reduction in PD-L1^+^ tumor cells and immune suppressive cell populations.

Changes in cytokine levels in the perfusate were also evaluated over time and in response to αPD-L1 (**Supplemental Fig. 5**). Levels of secreted anti-tumor interferon-γ (**Supplemental Fig. 5P**), increased following αPD-L1 treatment when compared to matched 3D-LTMs at day-6; no other significant changes were observed.

### Patient-Derived 3D Lung Tumor Models Simulate Response to αPD-1

Next, response to αPD-1 was evaluated, as described in **Figure 4A** in 3D-LTMs generated from 7 patient tumors (patient demographics found in **Table 2**). 3D-LTMs were considered responders if the difference in the ratio of PD-1^+^ CD8^+^ T cells to total CD8^+^ T cells or the differences in the ratio of PD-L1^+^ tumor cells to total tumors cells between patient matched IgG and αPD-1 treated 3D-LTMs was below −0.1, partial responders if the difference in the ratio of PD-1^+^ CD8^+^ T cells to total CD8^+^ T cells was negative but not below the −0.1 threshold, and non-responders if the difference in the ratio of PD-1^+^ CD8^+^ T cells to total CD8^+^ T cells was positive. Changes in cell viability were observed in responder (green), partial responder (purple) and non-responder (red) 3D-LTMs (**Fig. 4B-D**). The proportion of total immune cells (**Supplemental Fig. 6F**), CD8^+^ T cells (**Fig. 4E**) and PD-1^+^ CD8^+^ T cells (**Fig. 4F**) did not change in response in αPD-1 treatment in 3D-LTMs when all samples were combined (left), yet when samples were split by response status, responder 3D-LTMs had a 2.5 and 4.6 fold higher proportion of CD8 ^+^ T cells when compared to non-responder and partial responder 3D-LTMs, respectively (**Fig. 4E**, right). Additionally, a significant reduction in the proportion of PD-1^+^ CD8^+^ T cells was observed in responder 3D-LTMs (**Fig. 4F**, right). When the ratio of PD-1 ^+^ CD8^+^ T cells to total CD8^+^ T cells was calculated, 71% of 3D-LTMs had a decreased ratio indicating response or partial response to αPD-1 (**Fig. 4G**, left). When samples were split by responder status, a reduction in this ratio was observed only in responder and partial responder 3D-LTMs, whereas an increase in this ratio was observed in non-responder 3D-LTMs (**Fig. 4G**, right). Additionally, proliferating CD8^+^ T cells did not change in response to αPD-1 when all samples were combined (**Fig. 4H**, left), however this cell population was highest in partial responder 3D-LTMs and increased with αPD-1 (**Fig. 4H**, right). No difference in PD-L1^+^ tumor cells was observed with treatment (**Fig. 4I**, left), however this cell population was low in responder 3D-LTMs (**Fig. 4I**, right). When the ratio of PD-L1^+^ tumor cells to total tumor cells was calculated, 71% of 3D-LTMs had a reduced ratio indicating response or partial response to αPD-1 (**Fig. 4J**, left) and a reduction in this ratio was observed in only responder and partial responder 3D-LTMs (**Fig. 4J**, right).

**Figure 4:**
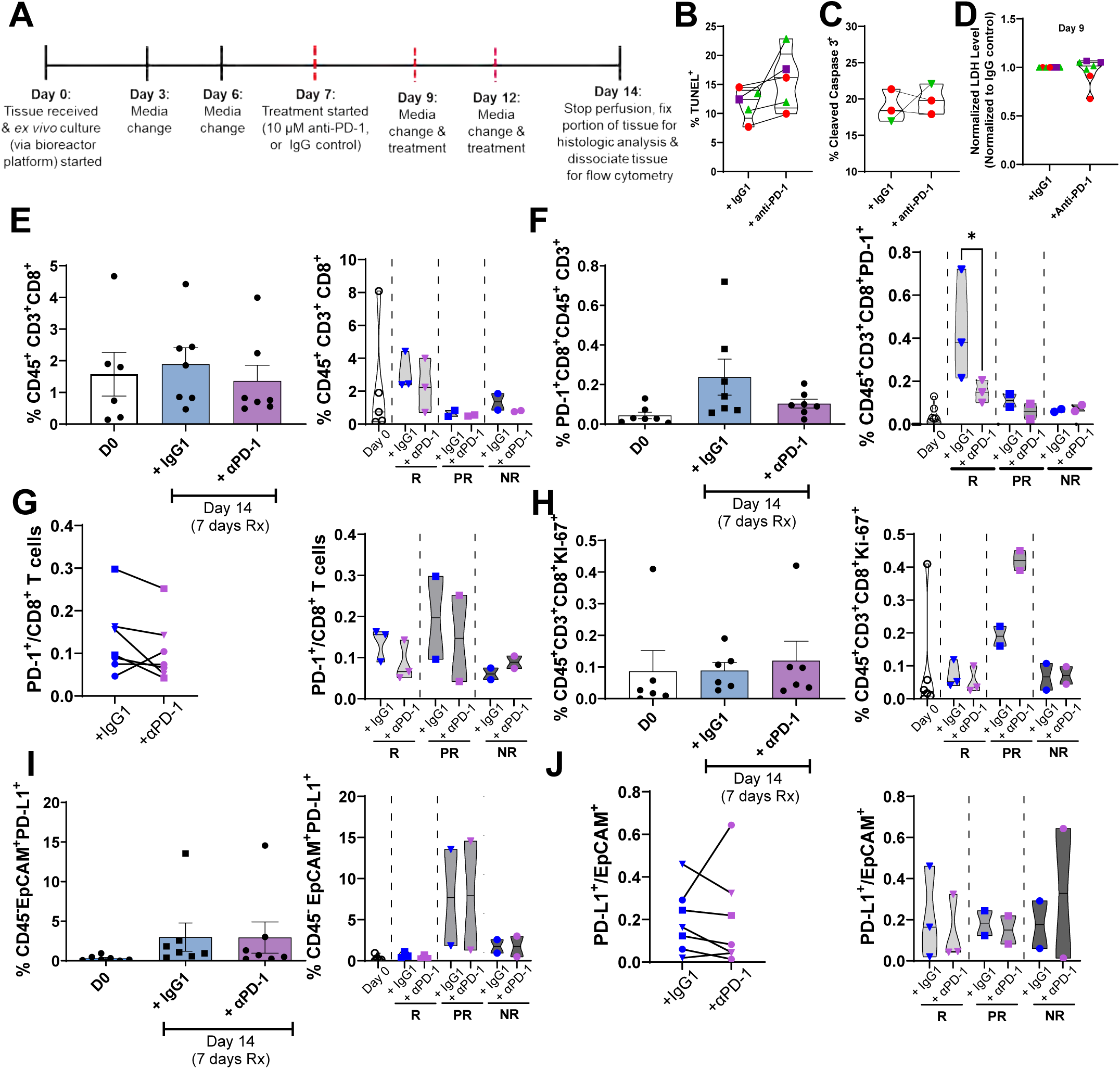
Patient-Derived 3D-LTM Model Response to αPD-1. **A.** 3D-LTM αPD-1 treatment schematic. **B-D.** Changes in cell viability are observed in responder (green), partial responder (purple) and non-responder (red) 3D-LTM. **E-F.** The proportion of CD8^+^ T cells (**E**) and PD-1^+^ CD8^+^ T cells (**F**) in 3D-LTM in response to αPD-1 or IgG treatment when all samples are combined (left) and when samples are split by response status (right). **G**. The ratio of PD-1 ^+^ CD8^+^ T cells to total CD8^+^ T cells is used to determine response status. **H.** Changes in the proportion of proliferating CD8^+^ T cells in 3D-LTM in response to αPD-1 or IgG treatment. **I**. The proportion of PD-L1^+^ tumor cells in 3D-LTM in response to αPD-1 or IgG treatment. **J**. The ratio of PD-L1^+^ tumor cells to total tumor cells is used to determine response status. * p<0.05.

Changes in tumor and stromal cells in 3D-LTMs in response to αPD-1 were also evaluated. Total tumor cells and hypoxic PD-L1^+^ tumor cells within 3D-LTMs did not change with αPD-1 (**Supplemental Fig. 6A-B**), when all samples were combined (left). When samples were analyzed based on responder status, total tumor cells tended to be lower in responder 3D-LTMs (**Supplemental Fig. 6A**, right) and hypoxic PD-L1^+^ tumor cells tended to be reduced in responder and partial responder 3D-LTMs (**Supplemental Fig. 6B**, right). Additionally, fibroblasts and PD-L1^+^ fibroblasts did not change with αPD-1 (**Supplemental Fig. 6C-D**, left), while endothelial cells tend to decrease with αPD-1 when all samples are combined (**Supplemental Fig. 6E**, left); this trend occurred across all response groups (**Supplemental Fig. 6E**, right). When the immune microenvironment was further assessed, the proportion of G-MDSC and M-MDSC did not change with αPD-1 when all samples are combined (**Supplemental Fig. 6F-H**, left).

Changes in cytokines secreted into the perfusate were also evaluated in response to αPD-1 (**Supplemental Fig. 7**). Among secreted cytokines, IL-4, IL-6, IL-17, and GM-CSF tended to be reduced in 3D-LTM conditioned media following αPD-1 when compared to matched isotype control treated samples (**Supplemental Fig. 7B, 7D, 7H, and 7N**). Additionally, significant reductions in IL-5 (**Supplemental Fig. 7C**) and G-CSF (**Supplemental Fig. 7O**) were observed following αPD-1 when compared to matched 3D-LTMs at day 6. IL-17 also tended to be reduced following αPD-1 when compared to matched 3D-LTMs at day 6 (**Supplemental Fig. 7H**). These results have been summarized in **Table 3**.

**Table 3:** Summary of 3D-LTM Response to ICI.

| Summary of Response: Split by Response Status |  |  |  |  |  |  |  |
| --- | --- | --- | --- | --- | --- | --- | --- |
|  |  | αPD-L1 |  |  | αPD-1 |  |  |
|  |  | Responder | Partial Responder | Non Responder | Responder | Partial Responder | Non Responder |
| Flow Cytometry | PD-L1+/ Total EpCam+ Tumor Cells | ↑ | ↓ | ↓ | ↓ | ↓ | ↑ |
|  | CD8+ T Cells | ↑ | No Change | ↓ | ↓ | No Change | ↓ |
|  | PD-1+/ total CD8+ T cells | ↓ | ↑ | ↑ | ↓ | ↓ | ↑ |
| Spatial Transcriptomics | Chemokine Signaling | N/A | Predominately increased | Mixed | Predominately increased | Predominately decreased | Little change |
|  | MHC Class II Antigen Presentation | N/A | Predominately increased | Predominately increased | Predominately increased | Predominately decreased | Some decreased |
|  | Dendritic Cell Activation | N/A | Predominately increased | Predominately decreased | Predominately increased | Predominately decreased | Little change |
|  | Mast Cells & IL-3 Signaling | N/A | Predominately increased | Predominately increased | Predominately increased | Predominately decreased | Little change |

### Differential Response to αPD-L1 and αPD-1 in 3D-LTMs

When changes in cell populations in response to αPD-L1 were compared to changes in response to αPD-1 variations were observed. The proportion of PD-L1^+^ tumor cells was reduced in response to αPD-L1 in responder and partial responder 3D-LTMs (**Fig. 3E**), whereas this cell population was low in 3D-LTMs that responded to αPD-1 (**Fig. 4I**). When the ratio of PD-L1^+^ tumor cells to total tumor cells was assessed, this metric was reduced in response to αPD-L1 in responder and partial responder 3D-LTMs but increased in non-responder 3D-LTMs (**Fig. 3F**), whereas this ratio was variable in response to αPD-1 (**Fig. 4J**). The proportion of CD8^+^ T cells increased in αPD-L1-responder 3D-LTMs but decreased in non-responder 3D-LTMs (**Fig. 3H**), whereas this cell population was high in 3D-LTMs that responded to αPD-1 (**Fig. 4E**). The proportion of PD-1^+^ CD8^+^ T cells decreased in αPD-L1-responder 3D-LTMs, but increased in non-responder 3D-LTMs (**Fig. 3I**). In αPD-1-responder 3D-LTMs, this cell population decreased and was low in non-responder 3D-LTMs (**Fig. 4F**). When the ratio of PD-1^+^ CD8^+^ T cells to CD8^+^ T tumor cells was assessed, this metric was reduced in response to both αPD-L1 (**Fig. 3J**) and αPD-1 (**Fig. 4G**) in responder 3D-LTMs, but increased in non-responder 3D-LTMs.

### Spatial Changes in Tumor-Immune Interactions in Response to αPD1

Next, spatial transcriptomics was completed to evaluate immune landscape changes within 3D-LTMs in response to ICI. Regions of interest (ROI) containing pan-cytokeratin^+^ tumor cells (green), CD45^+^ immune cells (red) and CD8^+^ T cells (yellow) were chosen as shown in **Figure 5A**. Differential gene expression was seen when ROIs from patient-matched IgG and αPD-1 treated 3D-LTMs (IgG: 3 ROI; αPD-1: 5 ROI) were compared (**Fig. 5B**), with 13 significantly upregulated and 32 significantly downregulated genes in αPD-1 treated samples. We next assessed changes in specific pathways between response groups. First, changes in the tumor inflammation signature (TIS) were assessed with treatment, as this gene signature has previously been associated with complete or partial response to αPD-1 treatment (34,35). ROI specific differences in the TIS within matched IgG and αPD1 treated responder 3D-LTMs were observed (**Fig. 5C**). We then compared TIS gene signatures (Log2 fold change (Log2FC)) relative to IgG control treatment, across the response groups following αPD-1 treatment, (**Fig. 5D**) and found that overall gene expression was highest in responder 3D-LTMs, indicating the presence of a more inflamed microenvironment in these tissues. Expression of CD8A and STAT1 was highest in responder 3D-LTMs, whereas TIGIT and CD27 were the only two genes upregulated in partial responders. All other genes in the TIS were downregulated, many significantly, following αPD-1 treatment, relative to IgG control in partial responder 3D-LTMs (Log2FC in **Supplemental Table 1**). Intermediate gene expression across the TIS was observed in non-responders. We then evaluated whether ICI modified expression of additional T cell immune checkpoints and found this pathway to be increased overall in responder 3D-LTMs, compared to partial and non-responders (**Fig. 5E**).

**Figure 5:**
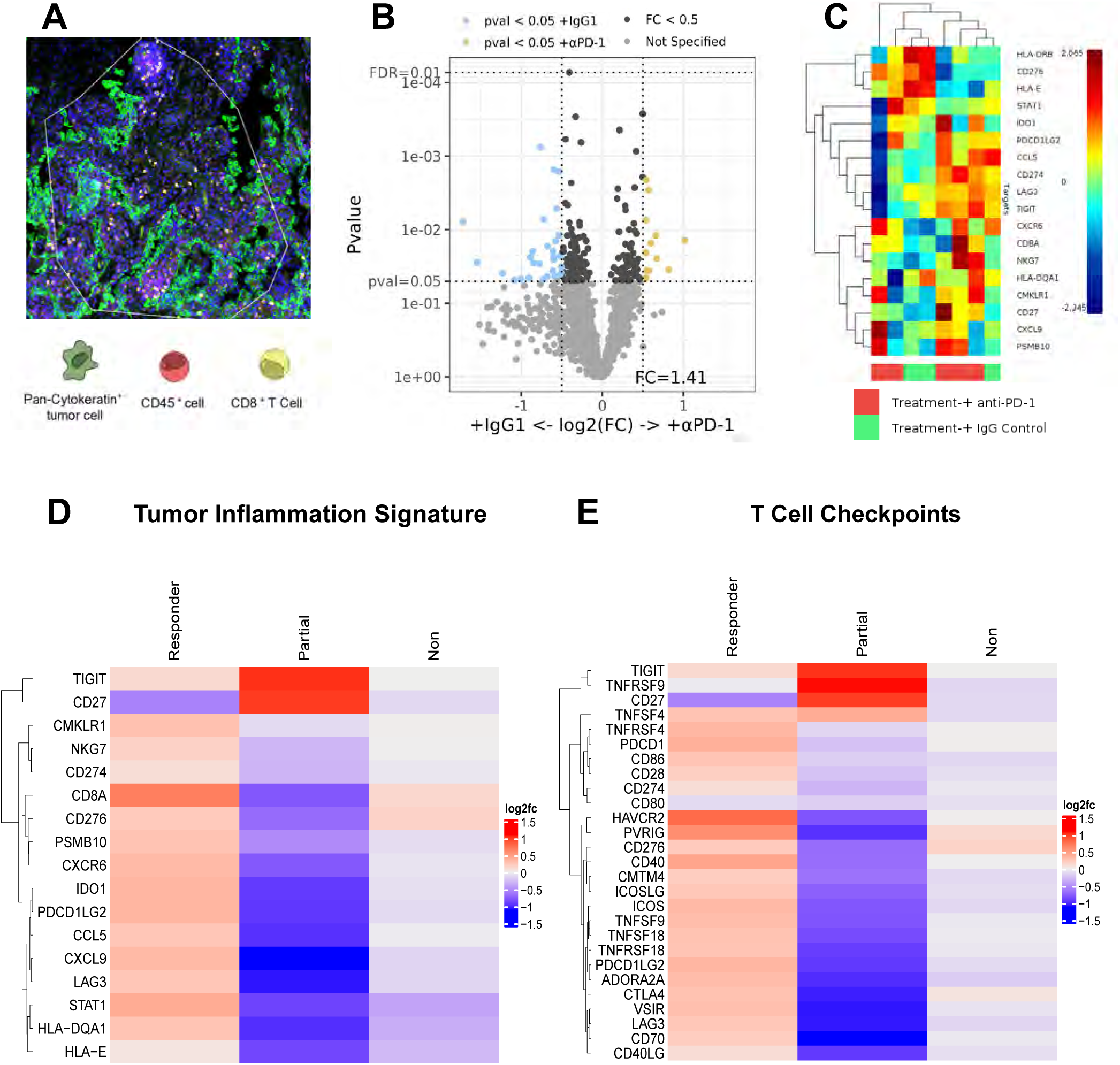
Spatial Changes in Tumor-Immune Interactions in Response to αPD1. **A.** Representative ROI showing pan-cytokeratin^+^ tumor cells (green), CD45^+^ immune cells (red) and CD8^+^ T cells (yellow). **B.** Volcano plot showing all gene expression differences between IgG and αPD1 treated responder 3D-LTM. **C.** Clustered heatmap with dendrogram showing ROI specific differences in the tumor inflammation signature (TIS) within matched IgG and αPD1 treated responder 3D-LTM. **D**. Heatmap comparing changes in TIS gene expression between responder, partial responder and non-responder 3D-LTMs. **E**. Heatmap comparing changes in T Cell Checkpoint gene expression between responder, partial responder and non-responder 3D-LTMs.

We then utilized GeoMx cell deconvolution algorithms to generate tissue maps displaying cell deconvolution proportions specific for each ROI in matched IgG and αPD1 treated 3D-LTMs (**Fig. 6A**). When cell proportions across samples were assessed, significant decreases in macrophages and CD4^+^ memory T cells and increases in CD8^+^ Memory T cells and dividing T cells, as well as an increasing trend in natural killer (NK) cell proportion were observed in response to αPD1 (**Fig. 6B**). Cell deconvolution showed that CD8^+^ memory T cells, plasmacytoid dendritic cells (pDCs), macrophages, classical monocytes, naïve CD4^+^ T cells, and plasma cells were increased in responder 3D-LTMs treated with αPD1 when compared to partial responder and non-responder 3D-LTMs (**Fig. 6C-E** and **Supplemental Fig. 8A-C**). Whereas, naïve CD8^+^ T cells, NK cells, neutrophils, naïve B cells, and CD4^+^ memory T cells were increased in non-responder 3D-LTMs when compared to partial responder and responder 3D-LTMs (**Fig. 6F-H** and **Supplemental Fig. 8D-E**). Additionally, endothelial cells, fibroblasts, non-classical monocytes, and regulatory T cells were increased in partial responder 3D-LTMs (**Supplemental Fig. 8F-I**). No change in the proportion of myeloid dendritic cells (mDCs) or memory B cells was observed and the average proportion of mast cells was similar across response groups (**Supplemental Fig. 8J-L**).

**Figure 6:**
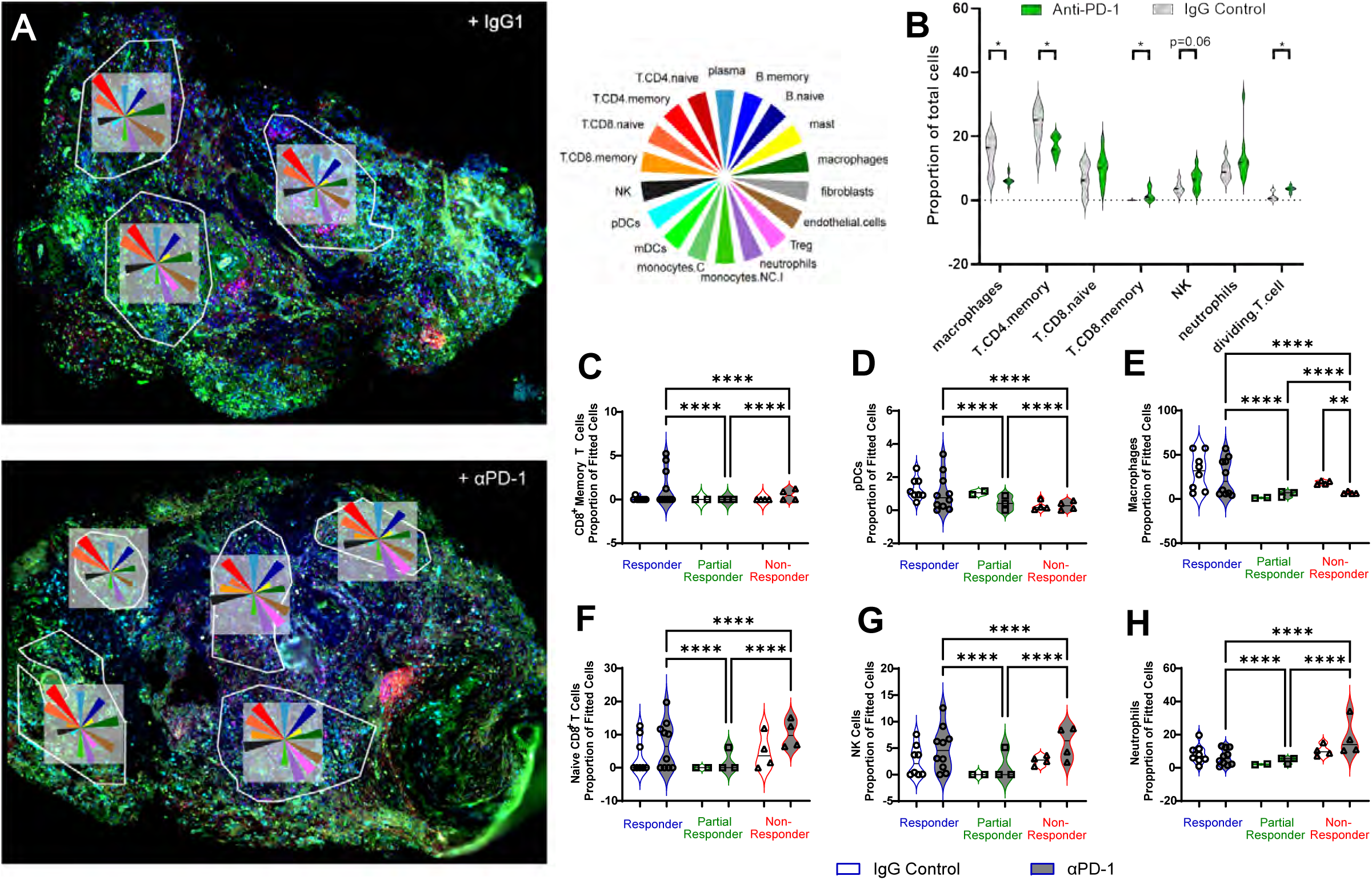
Spatial Changes in Cell Populations Response to αPD1. **A.** Representative tissue maps showing cell deconvolution proportions for specific ROI in matched IgG and αPD1 treated 3D-LTM. Wedge size is proportional to estimated cell counts. **B.** Changes in cell proportions in response to αPD1. **C-H.** Cell deconvolution showing proportion of CD8^+^ memory T cells (**C**), plasmacytoid dendritic cells (pDCs, **D**), macrophages (**E**), naïve CD8^+^ T cells (**F**), natural killer cells (**G**), and neutrophils (**H**) in ROIs of responder, partial responder, and non-responder 3D-LTMs following αPD1 or IgG treatment. * p<0.05; ** p<0.01; **** p<0.001.

When cell proportions were correlated with type of treatment (αPD1 or αPD-L1) or response status, mast cells, naïve B cells, CD4^+^ Memory T cells and endothelial cells were found to positively correlate with response status (**Supplemental Fig. 8M**). Plasma cells, CD8^+^ Memory and Naïve T cells, NK cells and non-classical monocytes showed a weaker, but still significant, positive correlation with response status and macrophages showed a negative correlation with response status (**Supplemental Fig. 8M**).

In Gene Set Enrichment (GSE) analyses, gene signatures differed in responder (**Fig. 7A**) and non-responder (**Fig. 7B**) 3D-LTMs treated with αPD-1, with activation in lymphocyte trafficking and chemokine response (green) noted in αPD-1 responder 3D-LTMs and suppression of antigen processing and presentation pathways (red) identified in non-responder 3D-LTMs. This was confirmed when gene signatures associated with chemokine signaling and MHC class II antigen presentation were compared across response groups (**Fig. 7C-D**). Heatmaps showed the highest gene expression in responder 3D-LTMs and low gene expression in non-responder 3D-LTMs. Significantly changing genes are shown in **Supplemental Table 1**. Chemokine signaling genes, CXCL11, GNG12, CCL15 (significantly upregulated when compared to IgG control, p=0.027), and CCL24 were the most upregulated in responder 3D-LTMs. GSK3B, GRB2, SHC3, CCL23, CCR9, PIK3CG, PRKACA, and STAT5B showed increased expression in partial responder 3D-LTMs. Whereas all other genes were decreased when compared to IgG control (57 of 121 significantly decreased, **Supplemental Table 1**), and showed relatively low expression when compared to other response groups. Non-responder 3D-LTMs show intermediate expression overall for most genes. Genes associated with MHC class II antigen presentation were relatively high in responder 3D-LTMs, particularly HLA related genes with HLA-DQB1 being the most upregulated when compared to IgG control (p=0.053), whereas partial responder 3D-LTMs have high expression of CTSL and reduced expression (8 of 21 significantly decreased, **Supplemental Table 1**) of all other genes. Non-responder 3D-LTMs have reduced expression of almost all genes in this pathway.

**Figure 7:**
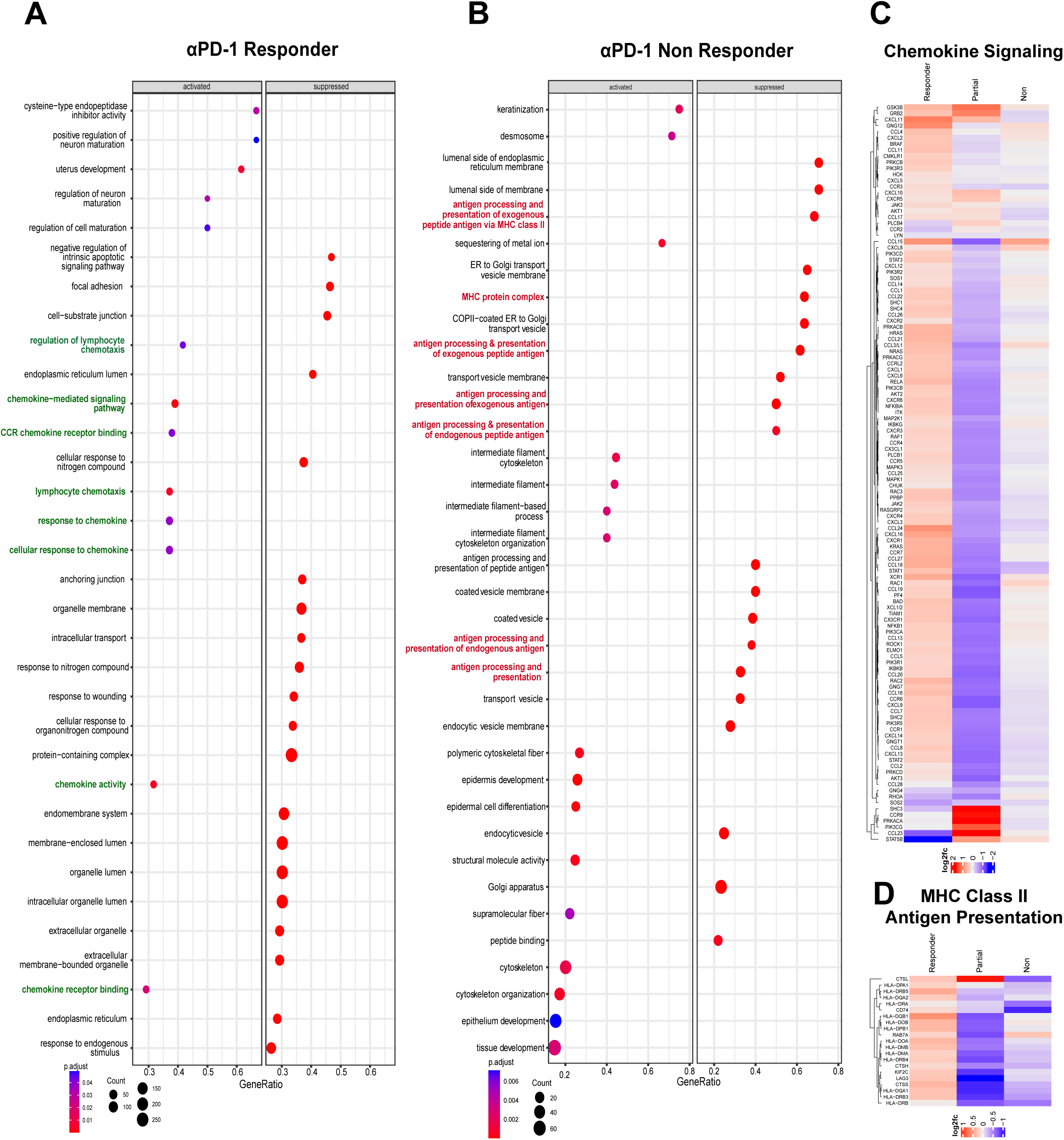
Gene Signatures Differ in Responder and Non-Responder 3D-LTM Suggesting Shifts in Lymphocyte Trafficking and Antigen Presentation. **A-B.** Pathway analysis shows activated lymphocyte chemotaxis and chemokine response and activity in 3D-LTM that respond to αPD-1 (**A**), whereas antigen processing and presentation are found to be suppressed in non-responder 3D-LTM (**B**). **C-G**. Heatmaps comparing gene expression changes in response to αPD-1 in Chemokine Signaling (**C**) MHC Class II Antigen Presentation (**D**), T cell Activation and Metabolism (**E**) Dendritic Cell Activation (**F**), and Mast Cell s& IL-3 Signaling (**G**) related genes between responder, partial responder and non-responder 3D-LTMs.

Additionally, gene expression associated with T cell activation and metabolism, dendritic cell activation, mast cells and IL-3 signaling were higher in responder 3D-LTMs and intermediate to low in non-responder 3D-LTMs. T cell activation and metabolism associated genes, including CD247, HAVCR2, CXCL11, and CD8A had relatively high expression and GIMAP6 had relatively low expression in responder 3D-LTMs. In partial responder 3D-LTMs only CXCL11, IL3, TIGIT, CD1C, STAT4, CXCL10, CXCR5, MLANA, DDP4, IL11 and CD27 showed increased expression when compared to IgG control. Most genes in the pathway had intermediate expression in non-responder 3D-LTMs (**Supplemental Fig. 9A**).

Genes associated with dendritic cell activation had relatively high expression in αPD-1 responder 3D-LTMs, with only LILRA4 and BATF3 showing relatively low expression. αPD-1 partial responder 3D-LTMs had low expression across the pathway with only ITGAX, CLEC5A, LILR4A having increased expression. Non-responder 3D-LTMs showed intermediate to low expression across all genes of the pathway. Similar results were observed when genes associated with mast cells and IL-3 signaling were assessed, with DDIT4 and FCER1G being the most upregulated in responder 3D-LTMs, IL3 and CSF2RB upregulated in partial responder 3D-LTMs and all other genes, including DDIT4 (trending reduction when compared to IgG control, p=0.14) and FCER1G (trending reduction when compared to IgG control, p=0.058) downregulated, and intermediate to low expression across the pathway in non-responder 3D-LTMs. This includes downregulation of DDIT4 (significantly reduced when compared to IgG control, p=0.034) and FCER1G (significantly reduced when compared to IgG control, p=0.022). These results are summarized in **Table 3**

When changes in cell proportions in response to αPD-L1 were assessed, partial responder 3D-LTMs were found to have higher proportions of CD8 populations, pDC, NK cells, neutrophils, endothelial cells, fibroblasts, mast cells, regulatory T cells, CD4^+^ memory T cells, and plasma cells (**Supplemental Fig. 10A-K**) and lower proportions of macrophages, monocytes populations, naïve B cells and naïve CD4^+^ T cells when compared to non-responder 3D-LTMs (**Supplemental Fig. 10L-P**). Little to no change in proportion of memory B cells or mDC was observed (**Supplemental Fig. 10Q-R**).

Pathway analysis on αPD-L1 treated 3D-LTMs showed that antigen presentation related pathways were suppressed in αPD-L1 partial responder 3D-LTMs and regulatory T cell differentiation related pathways were activated in non-responder 3D-LTMs (**Supplemental Fig. 11A-B**). Evaluation of chemokine signaling genes showed increased expression in partial responder 3D-LTMs when compared to non-responder 3D-LTMs (**Supplemental Fig. 11C**). T cell activation and metabolism genes also showed differential expression between response groups, with DPP4 and GIMAP6 increased in partial compared to non-responder 3D-LTMs (**Supplemental Fig. 11D**, as shown in **Supplemental Table 1**). Similar to what was observed in αPD-1 partial responder 3D-LTMs, TIS associated genes were relatively low, with only NKG7 (trending upregulation when compared to IgG control, p=0.13), TIGIT, and CD27 showing increased expression in αPD-L1 partial responder 3D-LTMs (**Supplemental Fig. 11E**). Furthermore, T cell checkpoint expression was increased in partial responder 3D-LTMs, with TIGIT, CD27, CD86, TNFSF4, TNFRSF9 showing the largest increase, similar to αPD-1 partial responders (**Supplemental Fig. 11F**). Genes associated with MHC class II antigen presentation had somewhat similar expression across response groups (**Supplemental Fig. 11G**). Unlike changes observed with αPD-1, dendritic cell activation related genes and genes associated with mast cells and IL-3 signaling showed generally higher expression in partial responder 3D-LTMs (**Supplemental Fig. 11H-I**). These results are summarized in **Table 3**.

In two patient samples, we evaluated response to both αPD-1 and αPD-L1 in 3D-LTMs. Based on ratios of PD-1^+^ CD8^+^ T cells to total CD8^+^ T cells and PD-L1^+^ tumor cells: total tumor cells, one was classified as a partial responder to both treatments (**Fig. 8A-B**). When gene signatures (normalized to IgG control) were compared between αPD-1 and αPD-L1 treated samples (Log2FC in **Supplemental Table 2**), overall expression within all pathways was higher with αPD-L1 treatment. Within TIS associated genes, reduced expression of CXCR6 (αPD-1 compared to IgG control, p=0.03), HLA-DQA1 (αPD-1 compared to IgG control, p=0.003), LAG3 (αPD-1 compared to IgG control, p=0.007), and NKG7 (αPD-1 compared to IgG control, p=0.13) was observed. While not significantly different from IgG control (p=0.13), NKG7 showed the largest increase in response to αPD-L1 (**Fig. 8C**). When genes associated with MHC class II antigen presentation were evaluated, only CTSL and CD74 were increased in αPD-1 treated 3D-LTM, whereas many HLA genes were significantly downregulated (**Fig. 8D**). All genes associated with chemokine signaling (**Fig. 8E**), except SHC3, PRKACA, and GRB2 were increased with αPD-L1 treatment. The largest reduction in dendritic cell activation genes in αPD-1 compared to IgG control were in BATF3 (p=0.018), SIGLEC5, SIGLEC8 (p=0.023), CD83 (p=0.001), and TLR7 (p=0.002; **Fig. 8F**). Genes associated with mast cells and IL3 signaling were also overall higher in αPD-L1 treated 3D-LTM, except for IL3, which was increased with αPD-1 treatment (**Fig. 8G**).

**Figure 8:**
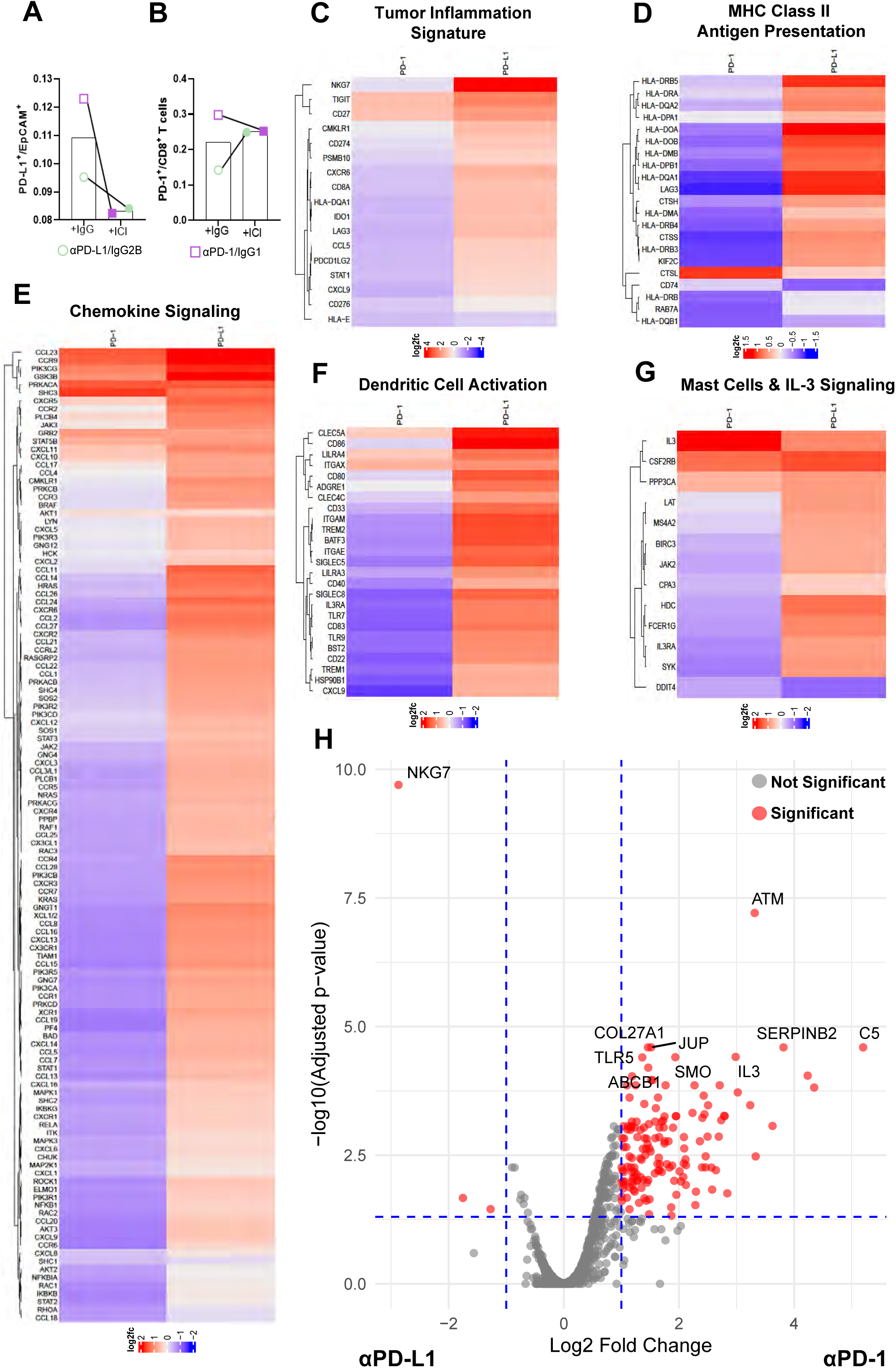
Gene Signatures Differ in Following αPD-1 and αPD-L1 in Partial Responder 3D-LTM from the Same Patient. **A-B**. Differences in the ratio of PD-L1 ^+^ tumor cells to total tumor cells (**A**) and the ratio of PD-1 ^+^ T cells to total CD8^+^ T cells (**B**) are observed in response to αPD-1 and αPD-L1 in 3D-LTM from the same patient, both 3D-LTM were considered partial responders. **C-G** Differential expression of TIS (**C**), MHC Class II Antigen Presentation (**D**), Chemokine Signaling (**E**), Dendritic Cell (**F**), and Mast Cells and IL-3 Signaling (**G**) related genes is observed in partial responder 3D-LTM generated from the same patient when response to αPD-1 and αPD-L1 is compared**. H**. Volcano plot showing differential gene expression in response to αPD-1 and αPD-L1 in 3D-LTM from the same patient.

Additionally, partial responder αPD-1 and αPD-L1 samples were compared via differential gene expression analysis and many genes were significantly upregulated in response to αPD-1 when compared to αPD-L1 treated 3D-LTM (**Fig. 8H**). This analysis showed that genes associated with immune regulation (i.e. KIR3DL3, INPP5D, TLR8, CD27 etc.) as well as genes associated with tumor suppression (i.e. SERPINB2, WNT7A, SHC3 etc.) were upregulated in αPD-1 treated 3D-LTM when compared to matched αPD-L1 treated 3D-LTM. NKG7 (p= 2E^-10^) and SLAMF6 (p=0.021), both involved in immune regulation, were the most upregulated gene in αPD-L1 treated 3D-LTM when compared to matched αPD-1 treated 3D-LTM. These results have been summarized in **Table 4**

**Table 4:** Summary Comparison of Tissue Matched 3D-LTM Response to ICI.

| Summary of Response: Tissue Matched Comparison |  |  |  |  |  |
| --- | --- | --- | --- | --- | --- |
|  |  | Tissue X |  | Tissue Y |  |
| | | αPD-L1<br>Partial<br>Responder | αPD-1<br>Partial<br>Responder | αPD-L1<br>Non<br>Responder | αPD-1<br>Responder |
| Flow Cytometry | PD-L1<sup>+</sup>/ Total<br>EpCam<sup>+</sup><br>Tumor Cells | ↓ | ↓ | ↑ | ↑ |
|  | CD8<sup>+</sup> T Cells | ↑ | ↑ | ↓ | ↑ |
|  | PD-1<sup>+</sup>/ total<br>CD8<sup>+</sup> T cells | ↑ | ↓ | ↑ | ↓ |
| Spatial<br>Transcriptomics | Chemokine<br>Signaling | Predominately<br>increased | Predominately<br>decreased | Predominately<br>increased | Predominately<br>increased |
|  | MHC Class II<br>Antigen<br>Presentation | Predominately<br>increased | Predominately<br>decreased | Predominately<br>increased | Mixed |
|  | Dendritic Cell<br>Activation | Predominately<br>increased | Predominately<br>decreased | Predominately<br>increased | Predominately<br>increased |
|  | Mast Cells &<br>IL-3 Signaling | Predominately<br>increased | Predominately<br>decreased | Predominately<br>increased | Predominately<br>increased |

The second patient 3D-LTMs tested with both αPD-1 and αPD-L1 was considered a responder to αPD-1 and a non-responder to αPD-L1 based on the ratios defined above (**Fig. 9A-B**). Consistent with previous observations, when evaluating TIS associated genes differential expression of TIGIT (p=0.12) and CD27 was observed with both genes increasing in responder αPD-1 treated 3D-LTM compared to non-responder αPD-L1 treated 3D-LTM (**Fig. 9C**). Expression of HLA-DRA, HLA-DMB, CD74, and HLA-DRB5, MHC class II antigen presentation associated genes, showed the largest increase (not significant) in responder αPD-1 treated 3D-LTM, whereas KIF2C (αPD-L1 compared to IgG control, p=0.017), and HLA-DRB4 (αPD-L1 compared to IgG control, p=0.006) were increased in non-responder αPD-L1 treated 3D-LTM (**Fig. 9D**). Chemokine signaling related genes showed relatively similar expression between responder αPD-1 treated 3D-LTM and non-responder αPD-L1 treated 3D-LTM, with the exception of CCL11 (significantly decreased when αPD-L1 compared to IgG control, p=0.04) and CCR9 (trending reduction when αPD-L1 compared to IgG control, p=0.08) which were increased in αPD-1 responder 3D-LTM and SOS2 which was increased in αPD-L1 non-responder 3D-LTM (**Fig 9E**). Dendritic cell activation associated genes TREM1 (αPD-L1 compared to IgG control, p=0.005) and F3 were increased in αPD-L1 non-responder 3D-LTM, whereas CLEC4C (αPD-L1 compared to IgG control, p=0.097), ITGAX, and CLEC5A were increased in αPD-1 responder 3D-LTM, (**Fig. 9F**). Among mast cell and IL-3 signaling associated genes, increased expression of DDIT4 (αPD-L1 compared to IgG control, p=0.03) and CSF2RB (responder αPD-1 treated 3D-LTM) was noted in responder αPD-1 treated 3D-LTMs. Whereas SYK (αPD-L1 compared to IgG control, p=0.003) and MS4A2 (trending increase when αPD-L1 compared to IgG control, p=0.12) were increased in non-responder αPD-L1 treated 3D-LTM (**Fig. 9G**). These pathway analyses could not fully explain the differential response observed with αPD-1 and αPD-L1 treatment in this patient sample, therefore responder αPD-1 and non-responder αPD-L1 3D-LTM were compared via differential gene expression analysis (**Fig. 9H**). This analysis showed that genes associated with immune regulation (TLR8, OLFML2B, CD79B etc.) and cell death (CASP1) were upregulated in αPD-1 treated responder 3D-LTM when compared to matched αPD-L1 treated non-responder 3D-LTM. NDUFA12 (p=0.003), and STAT5B (p=0.01), genes associated with tumor progression, and PRMT8 (p=0.004) were upregulated in αPD-L1 treated non responder 3D-LTM when compared to matched αPD-1 treated responder 3D-LTM. These results have been summarized in **Table 4**.

**Figure 9:**
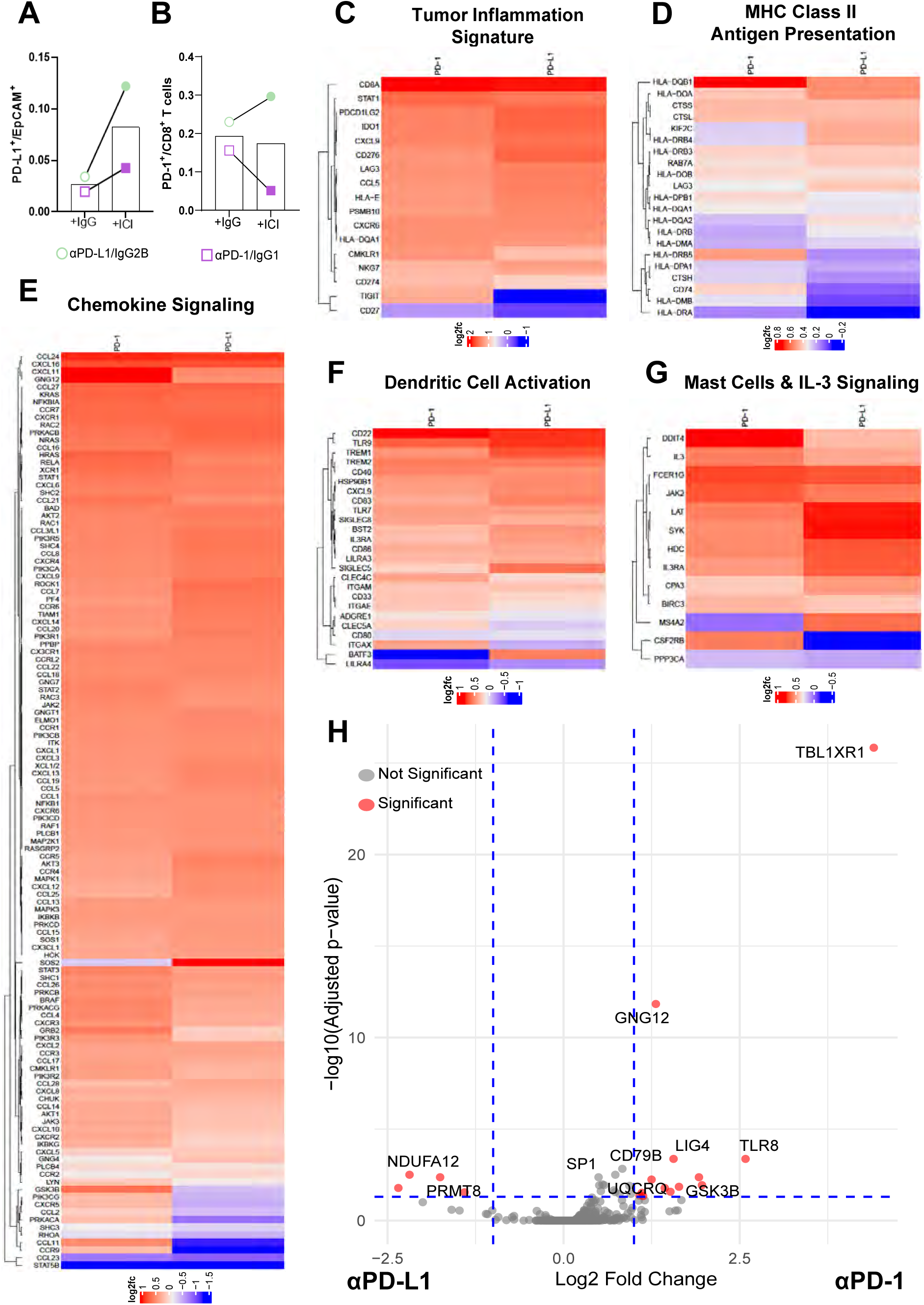
Gene Signatures Differ in αPD-1 Responder and αPD-L1 Non-Responder 3D-LTM from the Same Patient. **A-B**. Differences in the ratio of PD-L1 ^+^ tumor cells to total tumor cells (**A**) and the ratio of PD-1 ^+^ T cells to total CD8^+^ T cells (**B**) are observed in response to αPD-1 and αPD-L1 in 3D-LTM from the same patient. αPD-1 treated 3D-LTM responded to therapy and αPD-L1 treated 3D-LTM was considered a non-responder. **C-G** Differential expression of TIS (**C**), MHC Class II Antigen Presentation (**D**), Chemokine Signaling (**E**) Dendritic Cell (**F**), and Mast Cells and IL-3 signaling (**G**) related genes is observed in partial responder 3D-LTM generated from the same patient when response to αPD-1 and αPD-L1 is compared. **H**. Volcano plot showing differential gene expression in response to αPD-1 and αPD-L1 in 3D-LTM from the same patient.

## DISCUSSION

Herein we utilize 3D-LTMs to assess response to ICI and to determine the utility of this platform for future novel immune directed therapeutic testing. While ICI has transformed the treatment landscape for many cancers, only a fraction of NSCLC patients receives long-term benefit. It is estimated that ∼21-27% of patients receiving ICI alone or combined with another ICI as first-line therapy will experience primary resistance, defined as tumor progression within 6 months of ICI, with this number as high as 55% in patients with advanced disease (36–41). While the processes underlying ICI resistance are not fully understood, some mechanisms influencing primary resistance, including tumor intrinsic factors (lack of tumor immunogenicity, loss of tumor antigen or HLA expression and aberrant signaling) and extrinsic factors (presence of immune suppressive cell populations, T cell exhaustion and upregulation of alternative immune checkpoints, and altered metabolism) have been described (36,42–47) (48–50). Yet predictive biomarkers of resistance and clinical outcome are lacking (50).

Several prior studies have identified key human TME components, such as tumor-infiltrating lymphocytes (TILs) and tumor-associated macrophages (TAMs), as important determinants of response to immune checkpoint blockade. For example, Herbst *et. al*. demonstrated that a T-cell–inflamed phenotype characterized by high expression of PD-L1 and a Th1-type chemokine milieu was associated with response to anti–PD-1 therapy in NSCLC (51). Similarly, Thommen *et. al*. highlighted the prognostic and predictive relevance of intratumoral CD8^+^ T cells and exhausted T cell states, showing that a high density of these cell populations was associated with improved response to PD-1/PD-L1 inhibitors (52). Our study corroborates these findings by demonstrating that a high proportion of PD-1^+^ CD8^+^ T cells were associated with response to αPD-1. Importantly, in line with the observations by Romero *et. al*., our studies also highlight the potential role of chemokine signaling in response to ICI (53). Distinct from prior reports, our dataset allows integration of secreted proinflammatory cytokine signatures, transcriptomic profile and cell deconvolution, providing deeper insights into the context-specific role of the TME. For instance, while previous studies have largely focused on the presence of immune cells, our findings emphasize the functional status and spatial distribution of these populations, which may better reflect immune competence and therapeutic susceptibility.

Of the patients that initially respond to first-line ICI, ∼50-60% of patients will experience acquired resistance, where initial benefit from ICI (6 months or greater of stable disease, partial response or complete response) is followed by progressive disease (37,41,54–57). Mechanisms contributing to acquired resistance appear somewhat different than those contributing to primary resistance. To date, studies suggest defects in antigen presentation and IFN-γ signaling, alterations in tumor metabolism, and increased tumor hypoxia contribute to acquired resistance (56,58–60). In some studies, tumors with acquired resistance retain CD8^+^ T cells and have upregulation of IFN-γ, suggesting a persistent but insufficient anti-tumor immune response (56). Whereas other studies detect two patterns of acquired resistance; one with reduced tumor-infiltrating CD8^+^ T cells and PD-L1 expression after development of resistance, and another with high CD8^+^ T cell infiltration and PD-L1 expression, as well as elevated expression of other immune-inhibitory molecules (61).

Both primary and acquired resistance are a result of dynamic interactions between within the complex TME which are challenging to assess in human tumors. Our results show that 3D-LTMs mimic the heterogeneity in ICI response observed in patients. The responder 3D-LTMs have the highest CD8^+^ T cell populations, pDC, NK cells, and macrophages, intermediate abundance of neutrophils and mast cells, and low abundance of regulatory T cells. Whereas, partial responder 3D-LTMs have increased levels of T_regs_, intermediate levels of pDC, NK cells and reduced levels of neutrophils, and macrophages. Non-responder 3D-LTMs have reduced levels of pDC and mast cells but high levels of neutrophils. These alterations are similar to studies completed in patient tissues and preclinical models resistant to ICI, where T_regs_ and neutrophils were increased (62–67). In our study, pathway analysis shows chemokine signaling related pathways are activated in responder tissues (αPD-1), whereas antigen presentation related pathways are suppressed in partial (αPD-L1) and non-responder (αPD-1) tissues, and T_reg_ differentiation related pathways are activated in non-responders. This corroborates previous studies of ICI resistance, in which defects in antigen presentation and T_regs_ were associated with primary and acquired resistance (48,56,58,61). While here, we are assessing only one point in time, this model has the benefit of facilitating assessment of temporal changes in cytokine signatures and secreted factors. Additionally, for this study, spatial transcriptomic signatures were assessed for the entire ROI, therefore the gene expression is averaged across cell types within each ROI. In the future, segmentation will be utilized to better understand cell type specific changes in gene expression in response to intervention.

Preclinical testing remains a bottleneck in oncology drug development in part due to a lack of effective models that recapitulate the human TME. This is particularly important for the development and testing of immunotherapies. Currently, patient-derived organoids are widely utilized to mimic human tumors, yet these models generally lack immune and stromal cells and often have issues with normal lung epithelial outgrowth, in the context of lung cancer, making response to immune directed therapies challenging to assess. Like organoids, our patient-derived models can be generated from a variety of NSCLC histologic subtypes and tumor stages, yet our 3D-LTMs have the benefit of maintaining diverse cell populations within the TME while preserving the native tissue architecture and human stromal components. However, like all tumor models, our 3D-LTMs have limitations, including the requirement of freshly isolated tissue specimen for model generation and the current inability to propagate specific tissue models for future studies. Additionally, the long-term viability of 3D-LTMs beyond 14 days have to be evaluated, as we have not kept these models in long-term culture. Furthermore, our spatial profiling was limited by the available cell deconvolution algorithms which did not differentiate between certain cell subpopulations (CD8^+^ Memory subsets, myeloid-derived suppressor cells, macrophage subsets, etc.). Additional studies testing combinatorial strategies to increase efficacy of checkpoint inhibitors are underway using biosimilar immune checkpoint inhibitors and novel immune targeted therapies.

## ACKNOWLEDGEMENTS

The authors would like to acknowledge the UAB O’Neal Comprehensive Cancer Center (NIH CA013148), UAB Comprehensive Flow Cytometry Core Facility (NIH P30 AI27667, NIH CA013148), the UAB Tissue Biorepository, the UAB Pathology Core Research Laboratories, the UAB Mass Spectrometry/Proteomics (MSP) Shared Resource, the Emory Integrated Genomics Core, and Nanostring Technologies Technology Access Program for their help in data collection.

## ABBREVIATIONS

3D: Three-dimensional
3D-LTM: Three-dimensional Lung Tumor Model
D0: Day 0
D7: Day 7
D14: Day 14
DCC: Digital count conversion
DSP: Digital Spatial Profiling
ECM: Extracellular Matrix
ICI: Immune checkpoint inhibition
IRB: Institutional Review Board
LDH: Lactate dehydrogenase
MDSC: Myeloid-derived suppressor cells
M-MDSC: Monocytic myeloid-derived suppressor cells
NSCLC: Non-small Cell Lung Cancer
pDCs: Plasmacytoid dendritic cells
PDMS: Polydimethylsiloxane
PMN-MDSC: Polymorphonuclear myeloid-derived suppressor cells
ROIs: Regions of interest
TME: Tumor microenvironment

## DATA AVALIABILITY

The data generated in this study are available upon request from the corresponding author.

## AUTHOR CONTRIBUTIONS

K.F.G. was involved in bioreactor model design and set up, experimental design and execution, data collection, data analysis, and manuscript and figure preparation. S.S., S.L.S., S.S.D., and R.J.K. were involved in data collection, data analysis, and manuscript editing. K.P.H. was involved in bioinformatics analyses and manuscript editing. J.L.B. was involved in bioreactor design and production, and manuscript editing. M.K., S.P., Y.B., and A.D. were involved in experimental design and reviewing overall concepts and thought process of the manuscript. M.A. gave critical insights for the manuscript and was involved in manuscript editing. B.W. and J.M.D. facilitated collection of remnant surgical specimen and were involved in manuscript editing. J.S.D. was involved in experimental design, data interpretation, and manuscript and figure preparation.

## ETHICAL STATEMENT

This study was approved by the University of Alabama at Birmingham Institutional Review Board (IRB-300003092, IRB-300003384, IRB-300008998) and conducted following approved guidelines and regulations. Written informed consent was obtained from all participants with specimen collected under protocol IRB-300003384 and IRB-300008998. A waiver of consent was given for specimen collected under IRB-300003092.

