## Supplemental Figures for "Patient-Derived Three-Dimensional Lung Tumor Models to Evaluate Response to Immunotherapy"

### Supplemental Figure 1

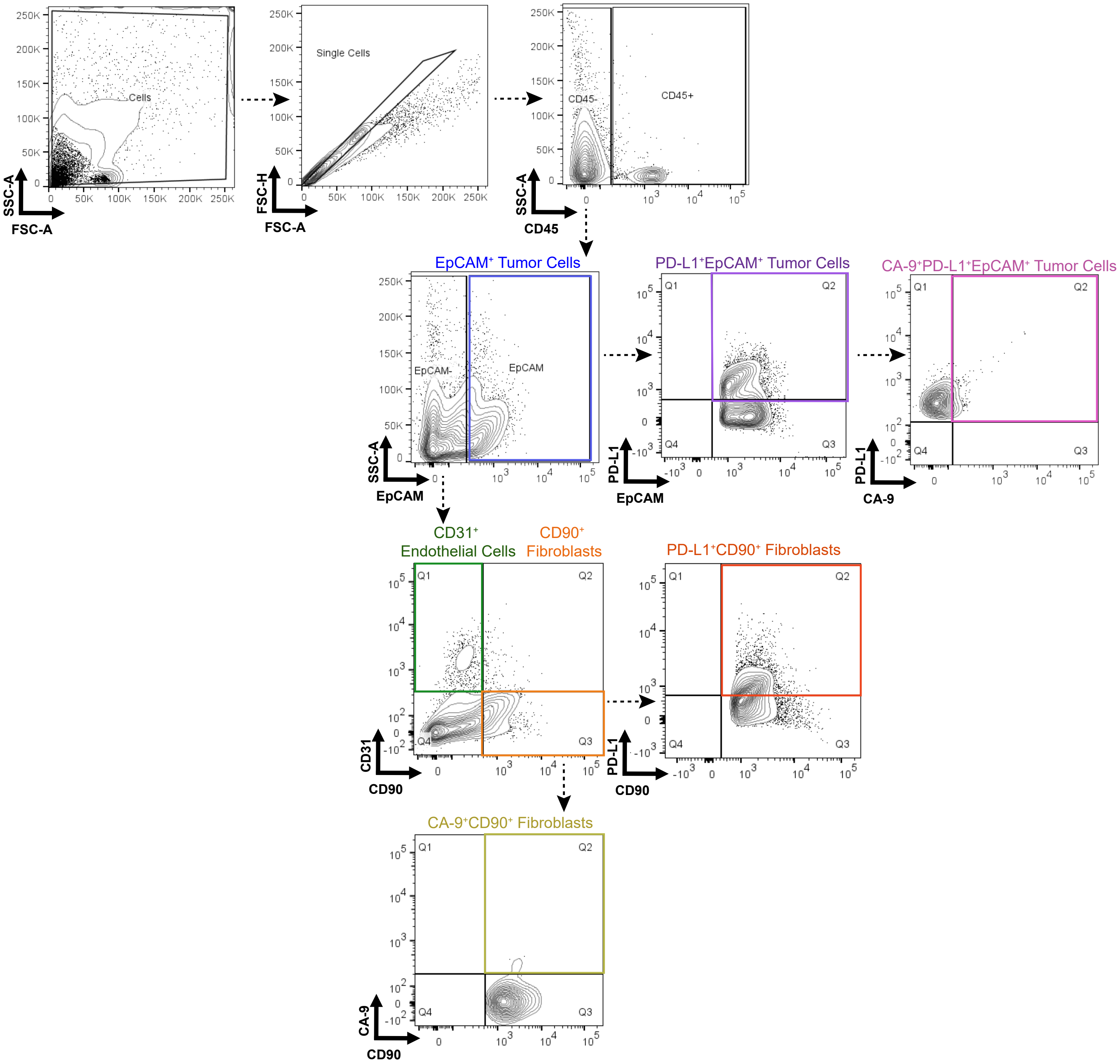

### Supplemental Figure 2

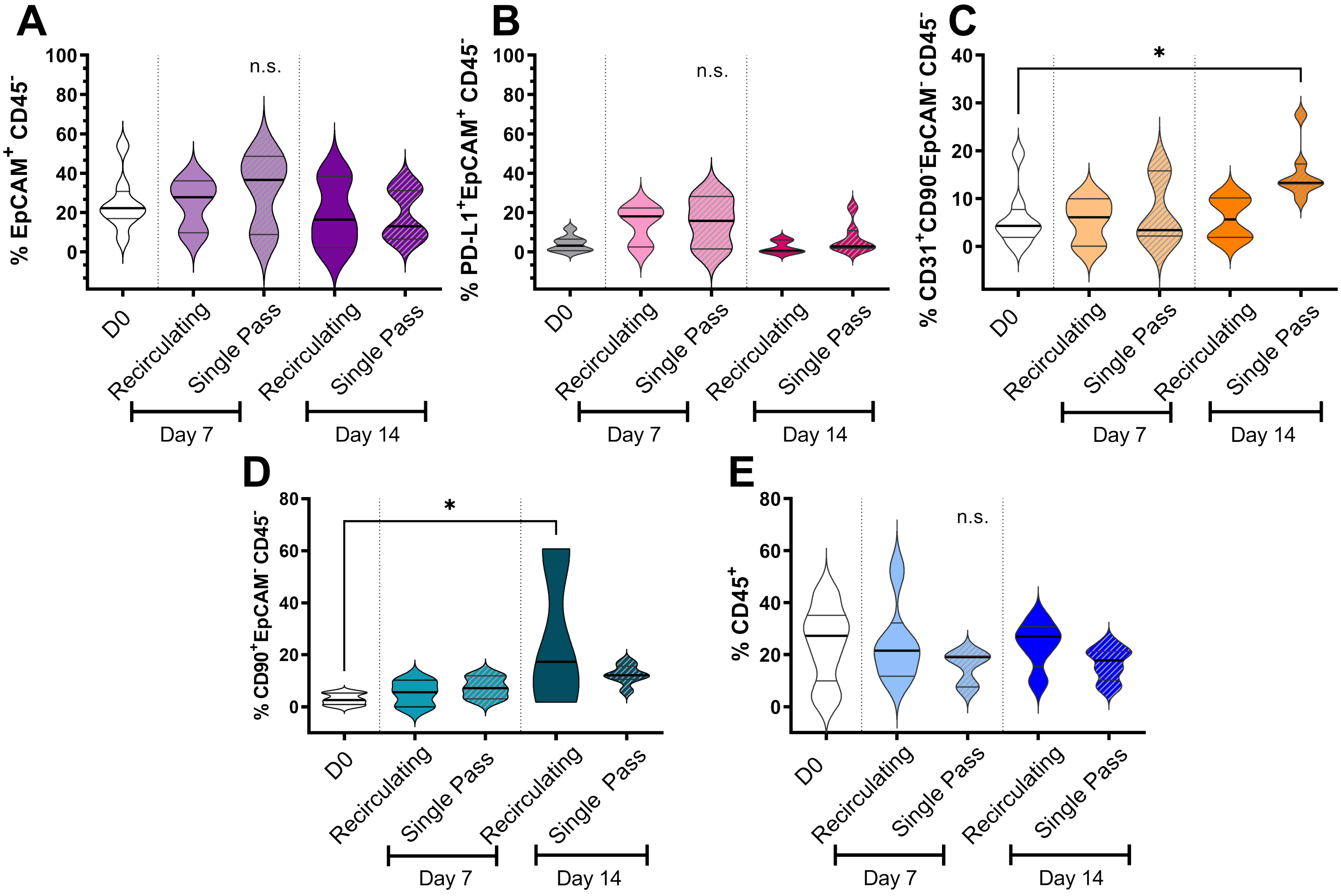

### Supplemental Figure 3

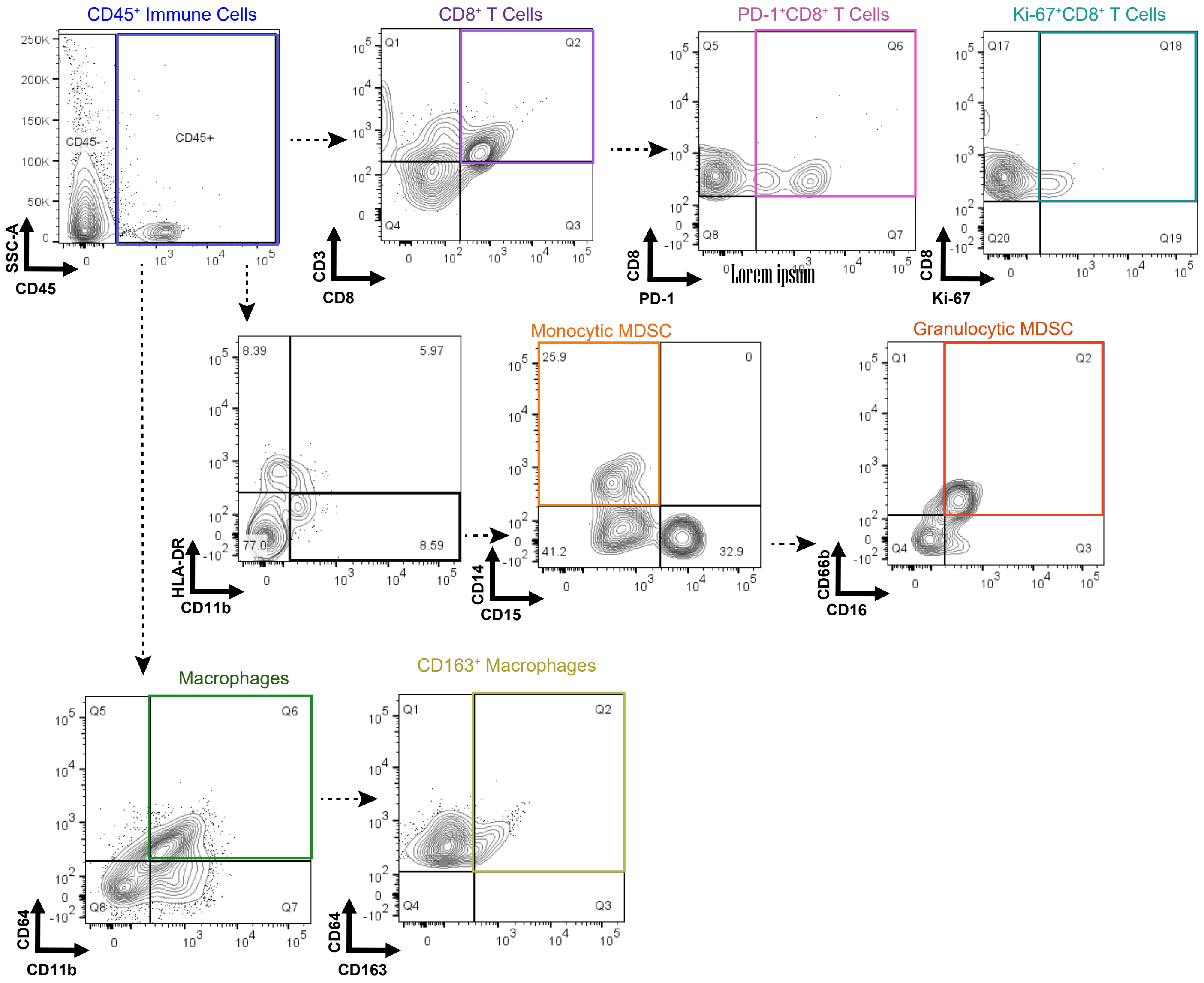

### Supplemental Figure 4

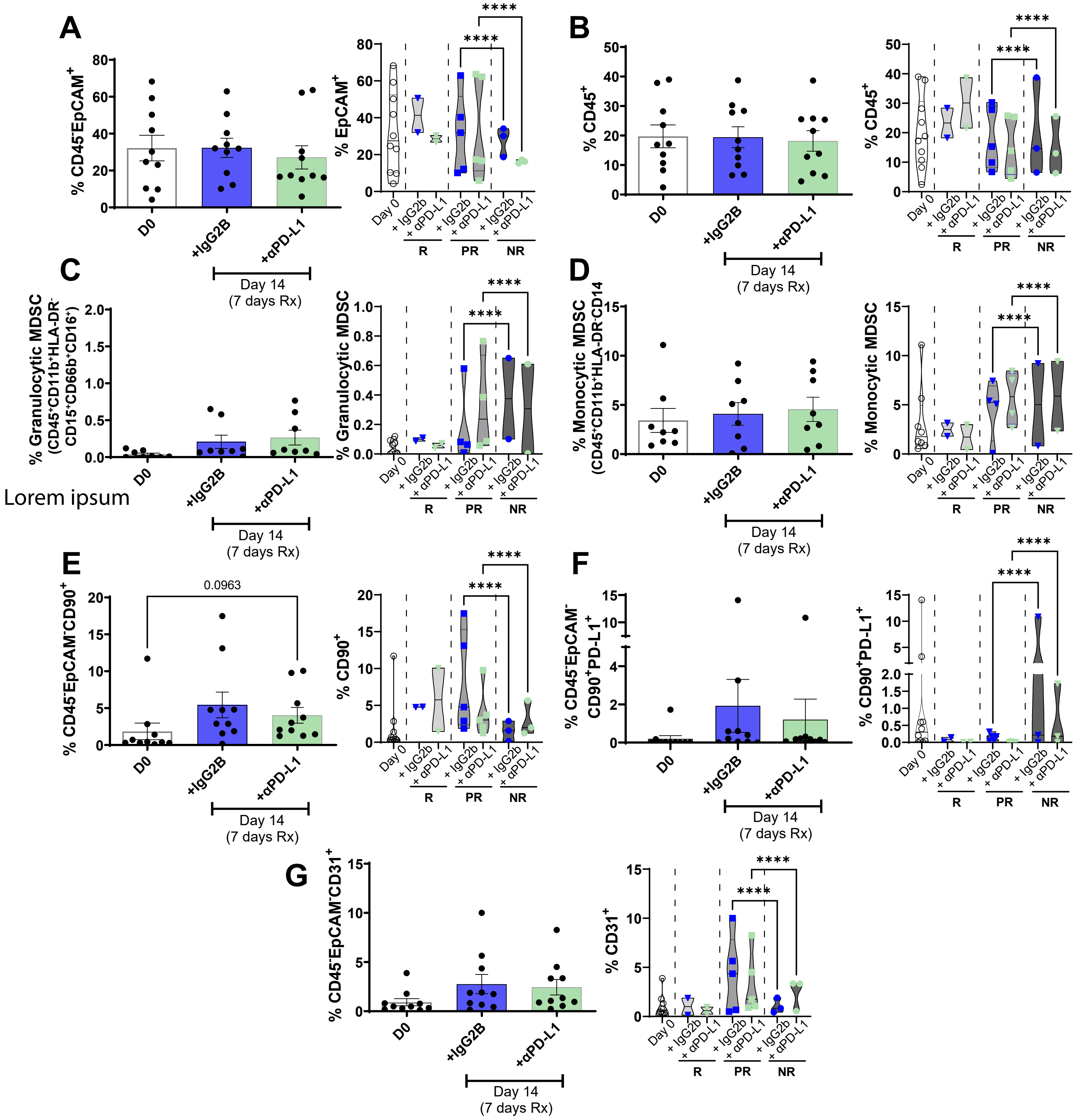

#### Supplemental Figure 5

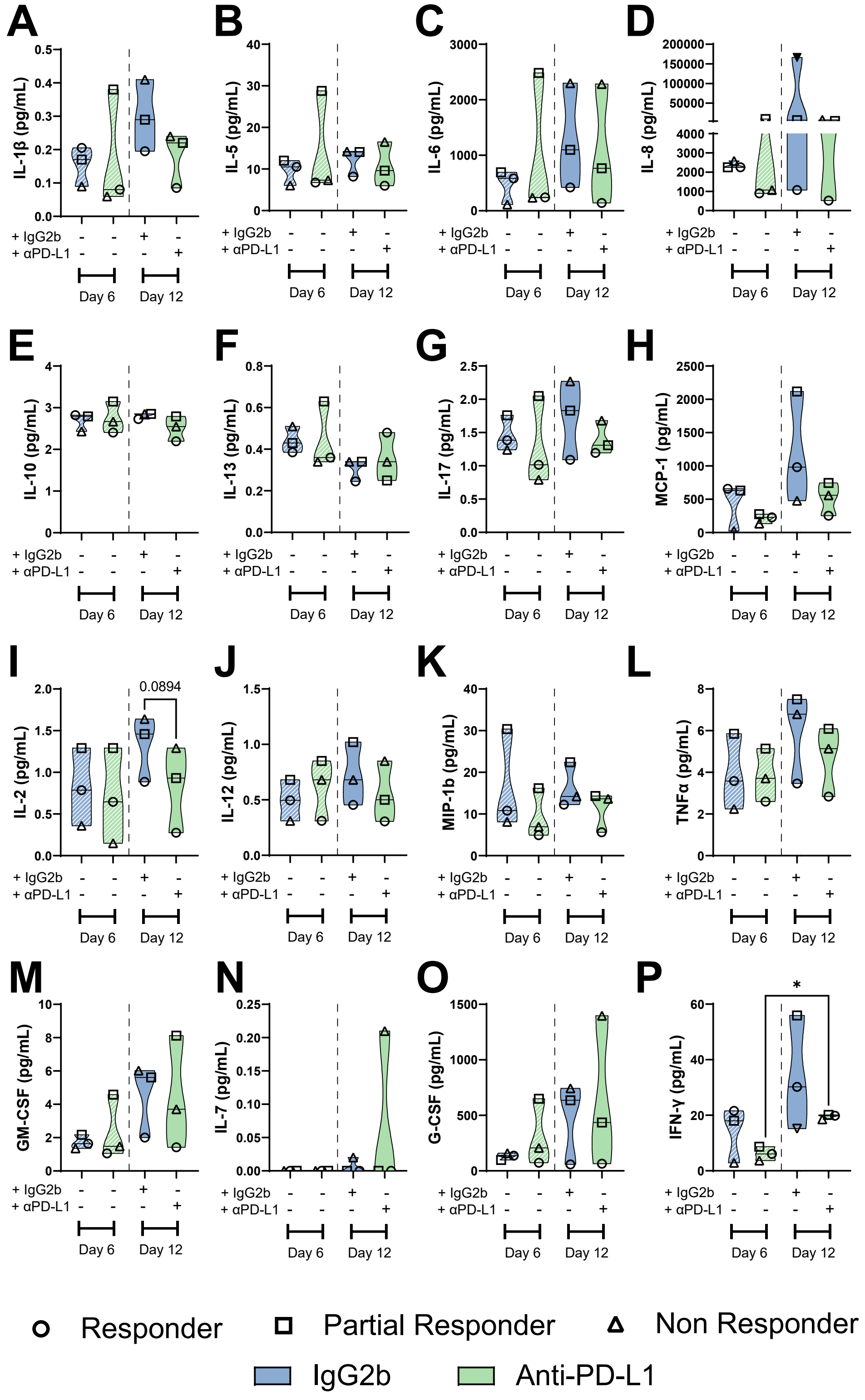

**Supplemental Figure 6**

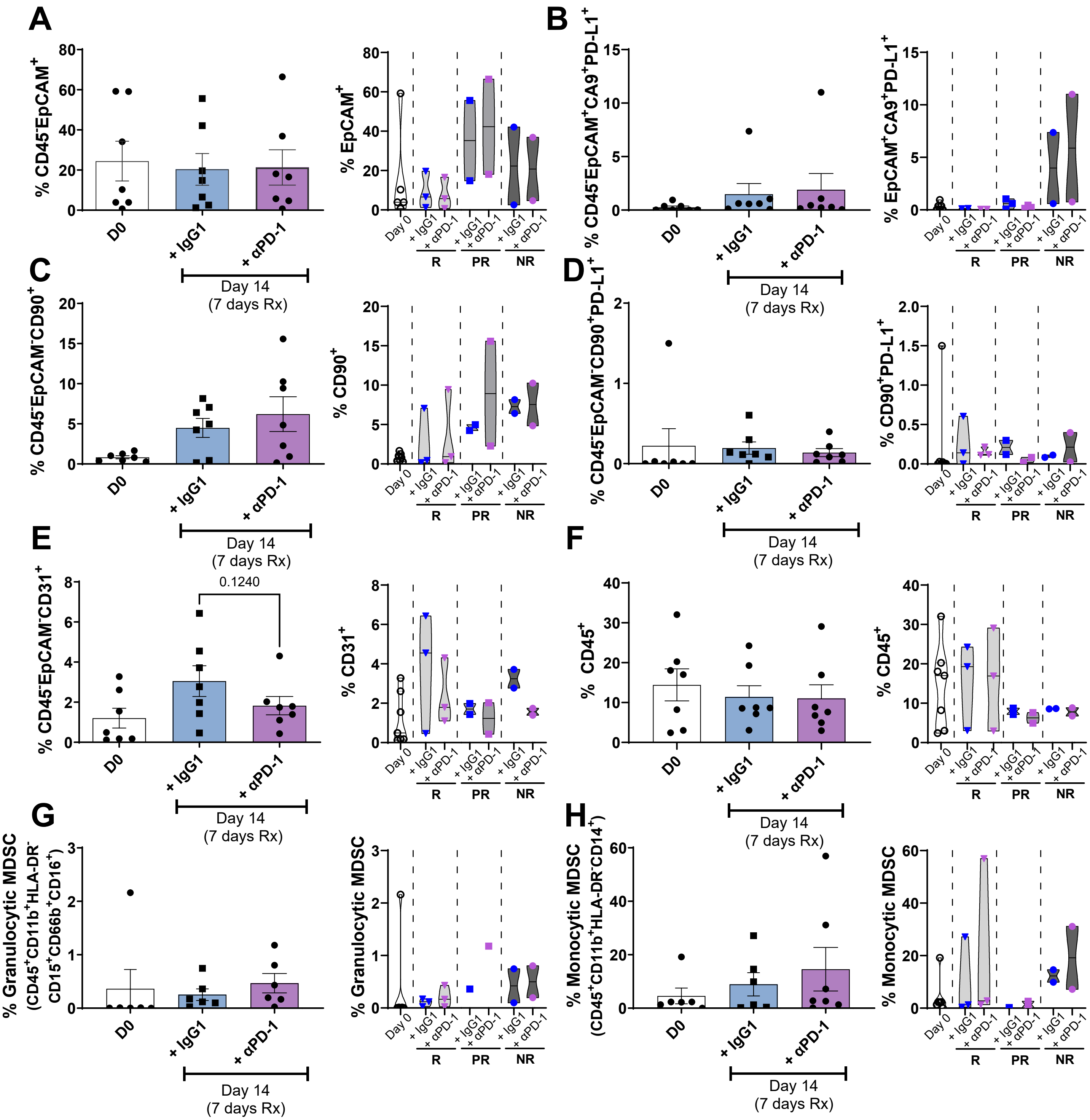

### Supplemental Figure 7

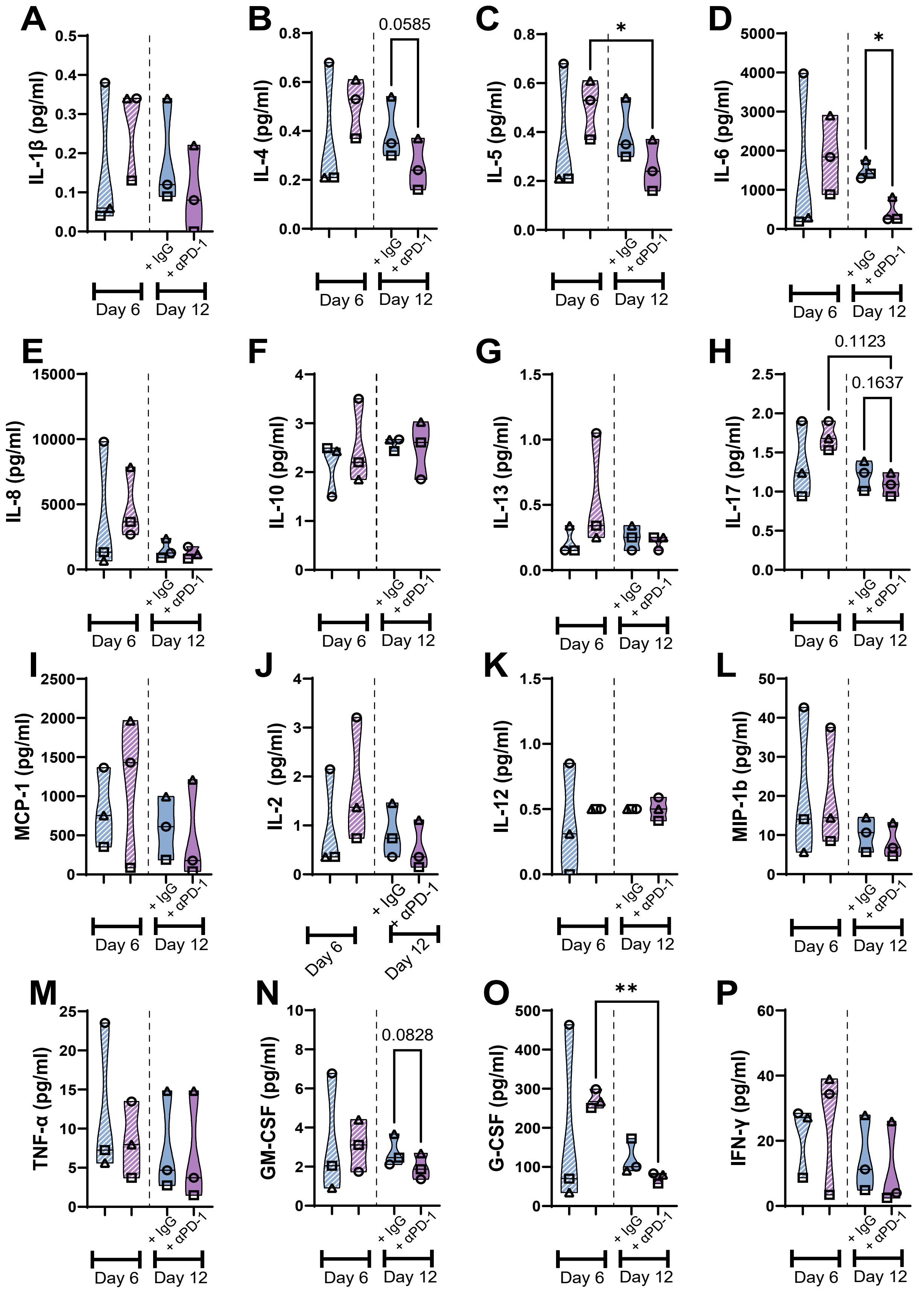

### Supplemental Figure 8

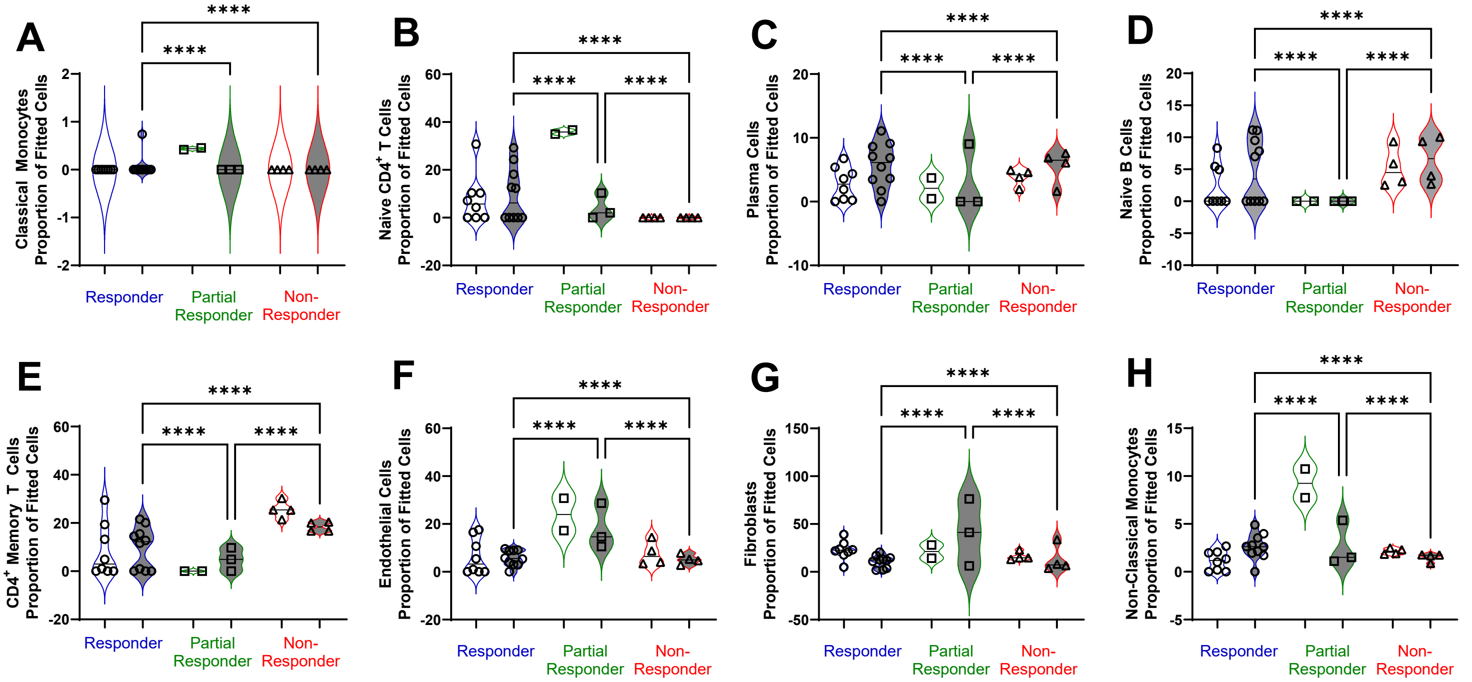


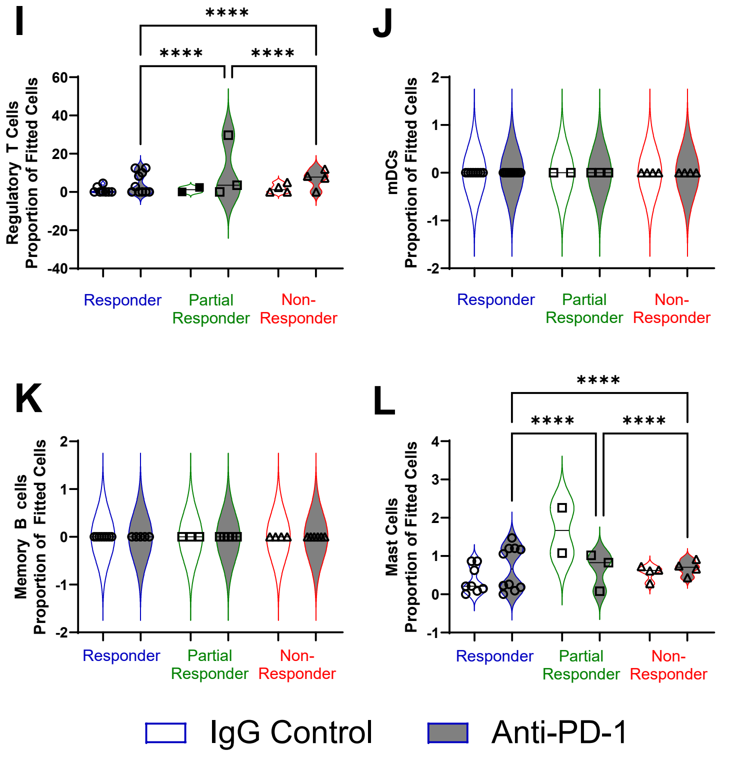


**M**

| | Correlation with Treatment | p value | Correlation with Response Status | p value |
|---|---|---|---|---|
| Macrophages | -0.102 | 0.583 | -0.437 | **0.014** |
| Mast Cells | -0.028 | 0.879 | 0.680 | **2.54E-05** |
| Naïve B Cells | 0.116 | 0.536 | 0.608 | **2.84E-04** |
| Memory B Cells | -0.201 | 0.278 | -0.155 | 0.405 |
| Plasma Cells | 0.083 | 0.655 | 0.391 | **0.030** |
| $CD4^+$ Naïve T Cells | -0.069 | 0.711 | -0.332 | 0.068 |
| $CD4^+$ Memory T Cells | 0.098 | 0.599 | 0.593 | **4.40E-04** |
| $CD8^+$ Naïve T Cells | 0.111 | 0.551 | 0.367 | **0.042** |
| $CD8^+$ Memory T Cells | 0.229 | 0.215 | 0.380 | **0.035** |
| NK | 0.146 | 0.433 | 0.392 | **0.029** |
| pDCs | -0.084 | 0.651 | -0.317 | 0.082 |
| Classical Monocytes | 0.163 | 0.382 | 0.325 | 0.075 |
| Non Classical Monocytes | 0.244 | 0.186 | 0.364 | **0.044** |
| Neutrophils | -0.268 | 0.144 | 0.129 | 0.489 |
| Treg | 0.266 | 0.148 | 0.255 | 0.166 |
| Endothelial Cells | 0.180 | 0.333 | 0.673 | **3.39E-05** |
| Fibroblasts | 0.014 | 0.940 | 0.190 | 0.305 |

### Supplemental Figure 9

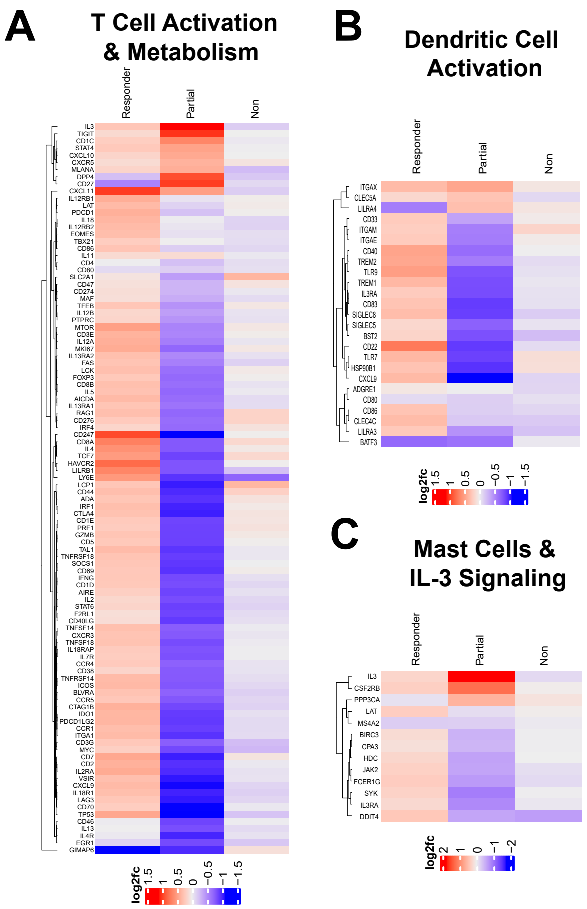

**Supplemental Figure 10**

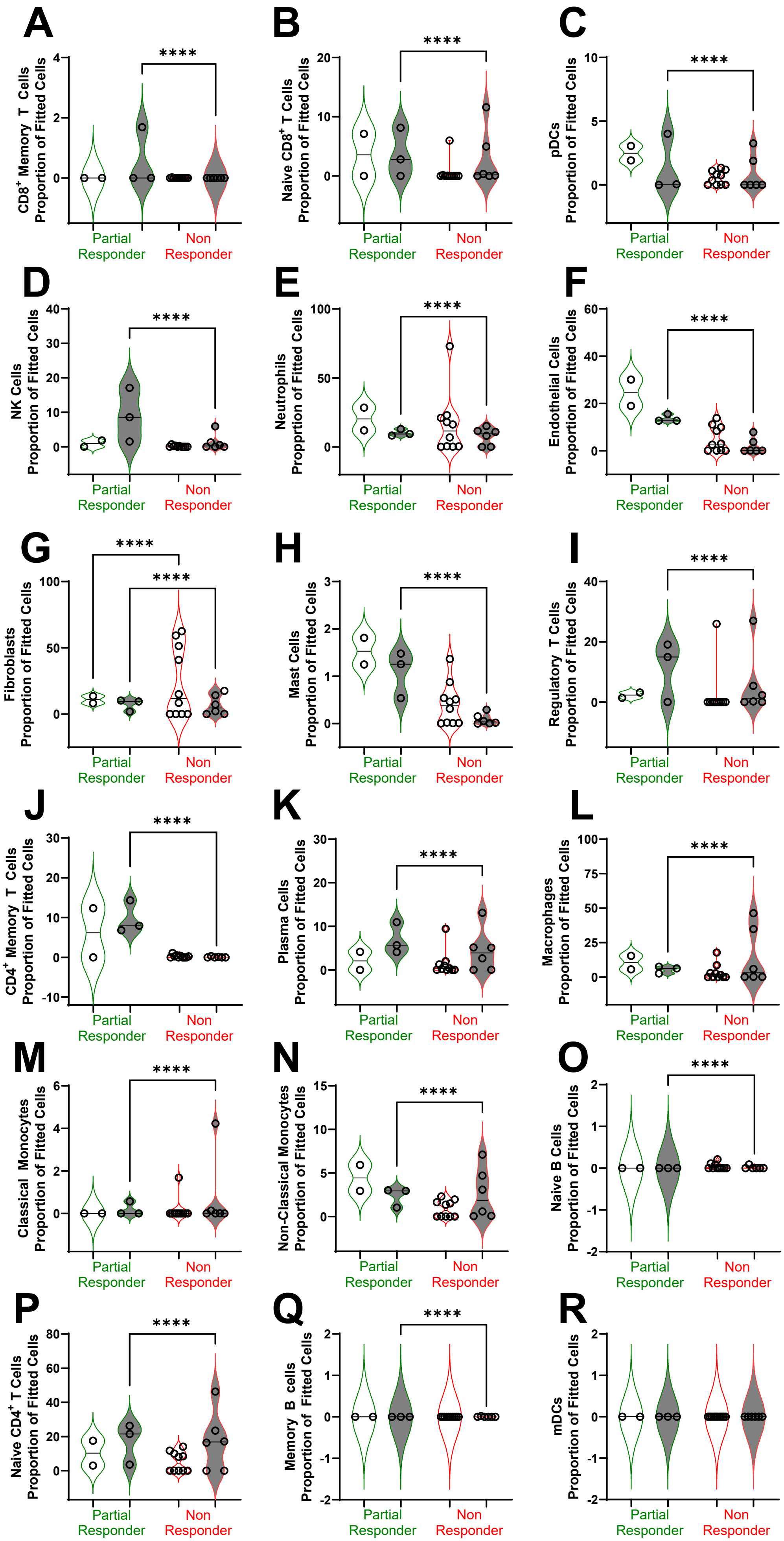

### Supplemental Figure 11

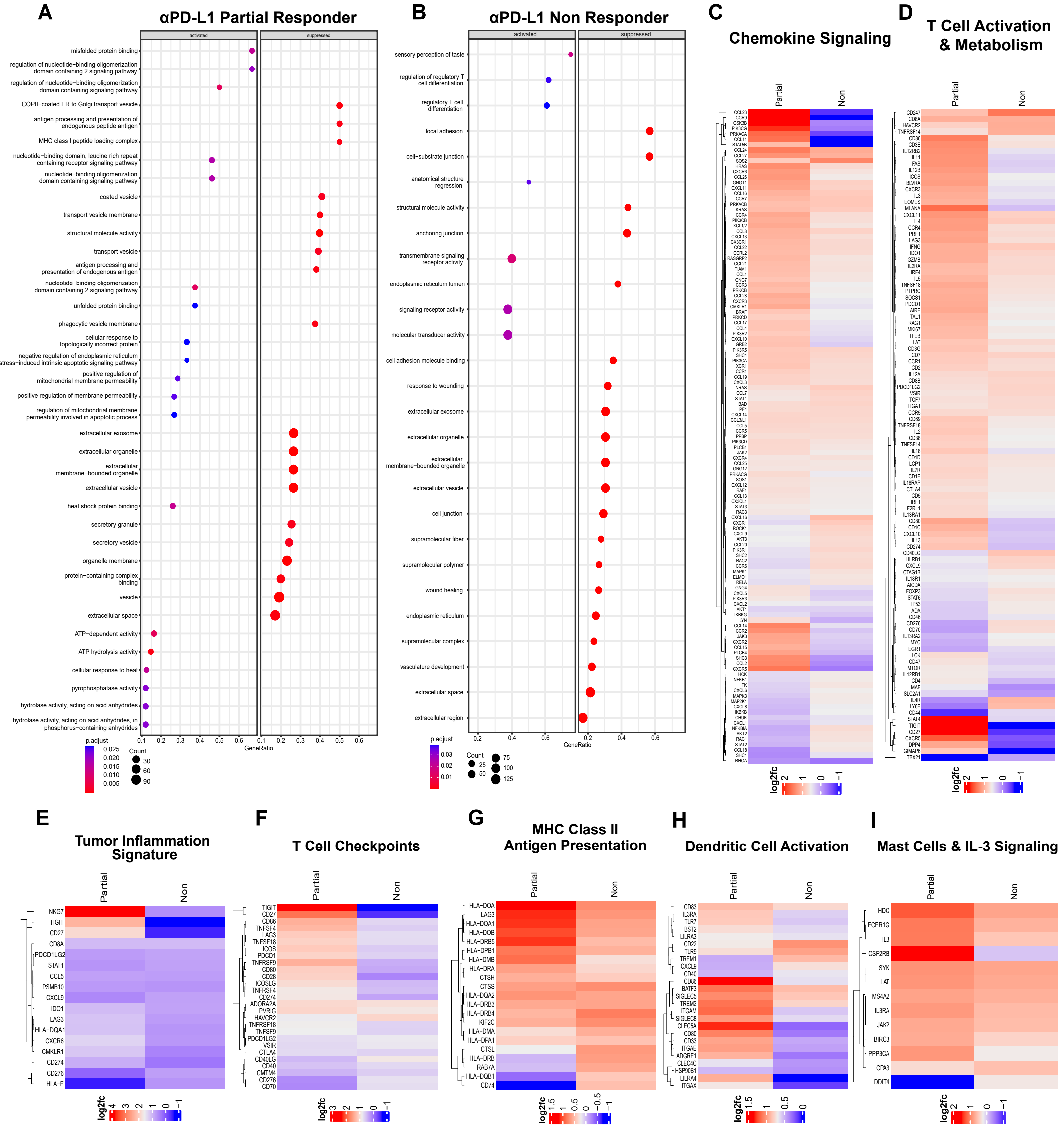
